# Polyalanine disease mutations impair UBA6-dependent ubiquitination

**DOI:** 10.1101/2022.06.20.496786

**Authors:** Fatima Amer-Sarsour, Daniel Falik, Yevgeny Berdichevsky, Alina Kordonsky, Gali Prag, Gad D Vatine, Avraham Ashkenazi

## Abstract

Expansion mutations in polyalanine stretches are now associated with a growing number of human diseases with common genotypes and similar phenotypes ^1–6^. These similarities prompted us to query the normal function of physiological polyalanine stretches, and investigate whether a common molecular mechanism is involved in these diseases. Here, we show that UBA6, an E1 ubiquitin-activating enzyme ^7, 8^, recognizes a polyalanine stretch within its cognate E2 ubiquitin-conjugating enzyme, USE1. Aberrations in this polyalanine stretch reduced ubiquitin transfer to USE1 and downstream target, the E3 ubiquitin ligase, E6AP. Intriguingly, we identified competition for the UBA6-USE1 interaction by various proteins with polyalanine expansion mutations in the disease state. In mouse primary neurons, the deleterious interactions of expanded polyalanine proteins with UBA6, alter the levels and ubiquitination-dependent degradation of E6AP, which in turn affected the levels of the synaptic protein, Arc. These effects could be observed in induced pluripotent stem cell-derived autonomic neurons from patients with polyalanine expansion mutations. Our results suggest a shared mechanism for such mutations, which may contribute to the congenital malformations seen in polyalanine diseases.

## Introduction

Trinucleotide repeats present within coding regions of the genome encode stretches of the same amino acid ^9^, and expansion mutations of such amino acid sequences have now been linked to a growing number of human diseases. The nine diseases caused by expansions of polyalanine stretches are due to the expression of aberrant nuclear proteins (eight of the nine are transcription factors). Examples include mutant paired like homeobox 2B (mutant PHOX2B) in congenital central hypoventilation syndrome (CCHS) ^1^, mutant RUNX family transcription factor 2 (mutant RUNX2) in cleidocranial dysplasia ^3^, mutant homeobox D13 (mutant HOXD13) in synpolydactyly^2^, and mutant poly(A) binding protein nuclear 1 (mutant PABPN1) in oculopharyngeal muscular dystrophy ^5^. The vast majority of cases are caused by *de novo* mutations and often involve congenital neurological abnormalities. The disease-causing expansions vary in length, according to the gene in question, with the severity of the associated clinical phenotype generally increasing with the length of the expanded tract ^9–11^. Physiological polyalanine stretches are commonly found in proteins, although they usually do not exceed 20 residues. In contrast, disease-causing expansion mutations may contain up to 13 additional alanine residues and can result in protein misfolding ^9–11^. The exact functional or structural role of the polyalanine stretches in normal proteins is unclear. One hypothesis is that they are simply protein spacer elements between functional domains. Another suggests that polyalanine stretches play roles in protein-protein recognition, but this remains unknown.

Perturbation of ubiquitin activation, conjugation, and transfer to target proteins via the E1-E2-E3 enzymatic cascade has been linked to dysregulation of neuronal homeostasis and to a growing number of neurodevelopmental disorders ^12–14^. Here, we describe a polyalanine motif in the ubiquitin-conjugating E2 enzyme UBE2Z/USE1, which contributes to USE1 ubiquitin loading by the E1 ubiquitin activating enzyme, UBA6. In addition, our results identify a domain in UBA6 that recognizes polyalanine-containing proteins and demonstrate that, under disease conditions, UBA6 preferentially interacts with different polyalanine-expanded proteins, thereby competing with USE1 binding. Since similar effects could be confirmed in neurons derived from patients with disease-causing mutations, we suggest that this represents a previously undescribed vulnerability caused by polyalanine expansion mutations.

## Results

A search for proteins containing polyalanine stretches in the ubiquitin cascades identified a small subset of E3 ubiquitin ligases and the E2 ubiquitin-conjugating enzyme, USE1 (**Extended Data Fig. 1a**). USE1 has both N- and C-terminal extensions on top of the ubiquitin-conjugating (UBC) core domain (**Fig. 1a**), classifying it as a class IV E2, which can be specifically loaded with ubiquitin by the E1 ubiquitin activating enzyme, UBA6, via a transthiolation reaction ^7, 8, 15^. The polyalanine stretch of USE1 is located in the N-terminal extension and is well conserved in primates, several other mammals, and reptiles (**Fig. 1a**, **Extended Data Fig. 1b**). In order to examine the role of the alanine stretch in USE1 ubiquitin loading, we replaced two alanine residues in the stretch with arginine residues. When transfected into HEK293T cells, the ubiquitin loading of the USE1 2A>2R mutant was significantly reduced in comparison to wild type USE1 (**Fig. 1b**). Exposure of cell lysates to the reducing agent β-mercaptoethanol, abolished USE1 loading under all examined conditions, suggesting that the additional band observed is ubiquitin conjugated via a thiolation reaction, similarly to a catalytic dead enzyme (USE1 C188A) that cannot be loaded with ubiquitin (**Fig. 1b**). Under non-reducing conditions, the ubiquitin loading of wild type USE1 and USE1 2A>2R was abolished in UBA6-siRNA depleted cells (**Fig. 1b**) suggesting that the difference in ubiquitin loading between the wild type and 2A>2R USE1 is dependent on UBA6 activity.

**Figure 1.**
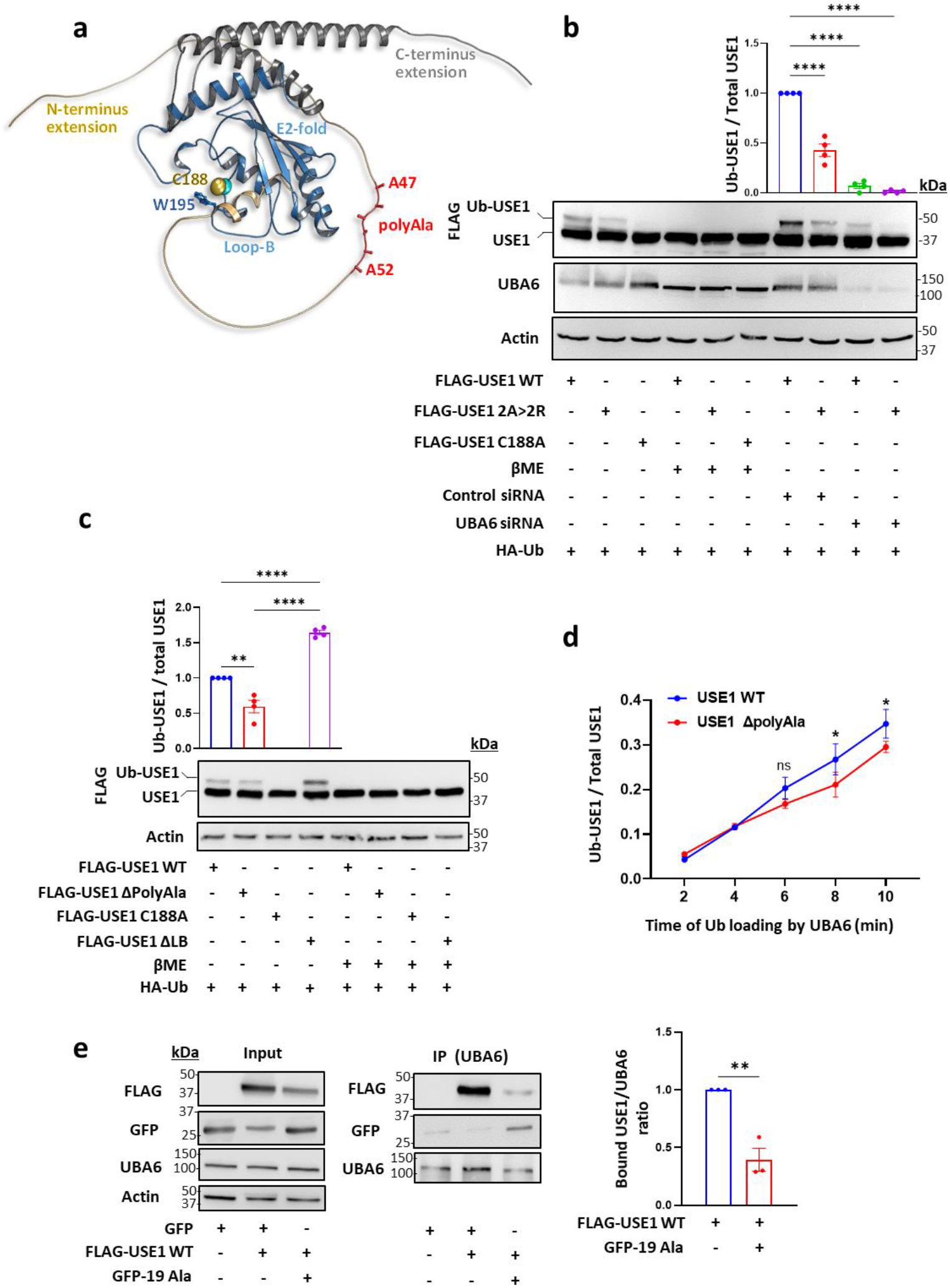
A polyalanine stretch in USE1 regulates UBA6-USE1 ubiquitin transfer. **a**, Structure of the USE1 enzyme (PDB 5A4P) indicating the catalytic cysteine (Cys188), loop B (LB) with Trp195 masking Cys188, and model of the C-and N-terminal extensions including the alanine repeats (created by AlphaFold). **b-c**, FLAG-wild type (WT) USE1, FLAG-USE1 mutants with aberrations in the polyalanine (2A> 2R, ΔPolyAla), FLAG-USE1 catalytic dead mutant (C188A), and FLAG-USE1 ΔLB were co-expressed with HA-Ub in control or UBA6-depleted (*Uba6* small inhibitory RNA, siRNA) HEK293T cells. Cell lysates were incubated with or without β mercaptoethanol (βME) and analyzed for ubiquitin loading. Results are mean ± s.e.m normalized to control WT USE1. One-way ANOVA Dunnet’s test (Fig.1b), Tukey’s test (Fig.1c), n=4. **d**, Time-dependent *in vitro* ubiquitin loading of WT and ΔPolyAla USE1 by UBA6. Quantification of USE1 ubiquitin loading is presented as mean ± s.e.m. Two-way ANOVA Sidak’s test, n=3. **e**, FLAG-WT USE1, empty GFP and GFP-polyAla (19Ala) constructs were transfected into HEK293T cells. Cell lysates were immunoprecipitated (IP) with anti-UBA6 antibodies and the immunocomplexes were analyzed with anti-FLAG, anti-UBA6 and anti-GFP antibodies. The bound USE1/UBA6 ratio is shown. Unpaired 2-tailed t-test, n=3. ns non-significant, * *P* < 0.05, ** *P* < 0.01, \*\*\*\**P* < 0.0001.

We constructed additional USE1 mutants including one with a deletion of the polyalanine stretch (USE1 ΔPolyAla), and a deletion of Loop B (USE1 ΔLB), which is hyperactive ^16^ (**Fig. 1c**). The USE1 ΔPolyAla mutant showed reduced ubiquitin loading compared to wild type USE1 or to USE1 ΔLB both in cells and *in vitro* with purified recombinant UBA6 (**Fig. 1c**, **Fig. 1d**). These findings provide further support that the polyalanine stretch contributes to the recognition of USE1 by UBA6.

### Isolated polyalanine stretches compete with USE1

In order to examine whether isolated polyalanine stretches (19 Ala residues) interact with UBA6 in mammalian cells, we transfected HEK293T cells with GFP-polyAla, and monitored the binding to UBA6, or its possible competition with USE1. We found that the polyAla stretch binds UBA6 and that this interaction decreases the USE1-UBA6 binding (**Fig. 1e**). The ubiquitin-activating E1 enzymes, UBA1 and UBA6 share 40% identity of protein sequence and a strong specificity for their cognate ubiquitin-like proteins ^17–19^. The ubiquitin fold domain of E1 interacts with the α1 helix of E2 and functions to determine selectivity between E2s of different ubiquitin-like proteins ^20^. To identify regions in UBA6 that are likely to determine the specificity to USE1, we compared the physico-chemical properties of UBA6 to those of UBA1 by structural modeling (**Fig. 2a**). Interestingly, comparison of the structures of UBA1 and UBA6 at the interface with the E2-α1 as well as the E2-α1 helices themselves seemed to be very similar. However, projection of calculated electro-potential properties on the surfaces of the two enzymes revealed significant differences within the second catalytic cysteine half (SCCH) domains. In place of the negative groove seen in UBA1, UBA6 contains a large positively charged groove that is formed by the key residues Lys628, Arg691, Lys709, and Lys704 (**Fig. 2a**). The structural comparison of the canonical E1 enzymes UBA1, 2, 3, 6 and 7 ^19–22^ shows significant differences in the SCCH domains (**Extended Data Fig. 2**). Moreover, a recent study demonstrated that replacing the SCCH domain of UBA1 with the one of UBA2 allows the engineered UBA1 to load ubiquitin onto the E2 of SUMO (UBC9) ^23^. Consequently, we assumed that the SCCH domain of UBA6 may hold specificity to USE1 and thus mutating the above-mentioned positive charged residues in UBA6 may affect the binding to USE1. Based on this hypothesis, we constructed UBA6 mutants with the positive residues replaced by Ala (UBA6 mut 4Ala) or Asp (UBA6 mut 4Asp). The USE1 binding of both these mutants was indeed significantly reduced in HEK293T cells (**Fig. 2b**) as was the interaction of both mutants with isolated polyalanine stretches compared to wild-type UBA6 (**Fig. 2c**). *In vitro*, both recombinant UBA6 mut 4Ala and UBA6 mut 4Asp showed reduced ability to load USE1 with ubiquitin when compared to wild type UBA6 (**Fig. 2d**). Interestingly, neither mutations in the positive groove of UBA6 nor aberrations in the polyAla sequence in USE1 are sufficient to abrogate ubiquitin transfer from UBA6 to USE1. This suggests that while these two regions are important for specificity they are not essential for the general mechanism of ubiquitin activation (*i.e*. adenylation) or transthiolation.

**Figure 2.**
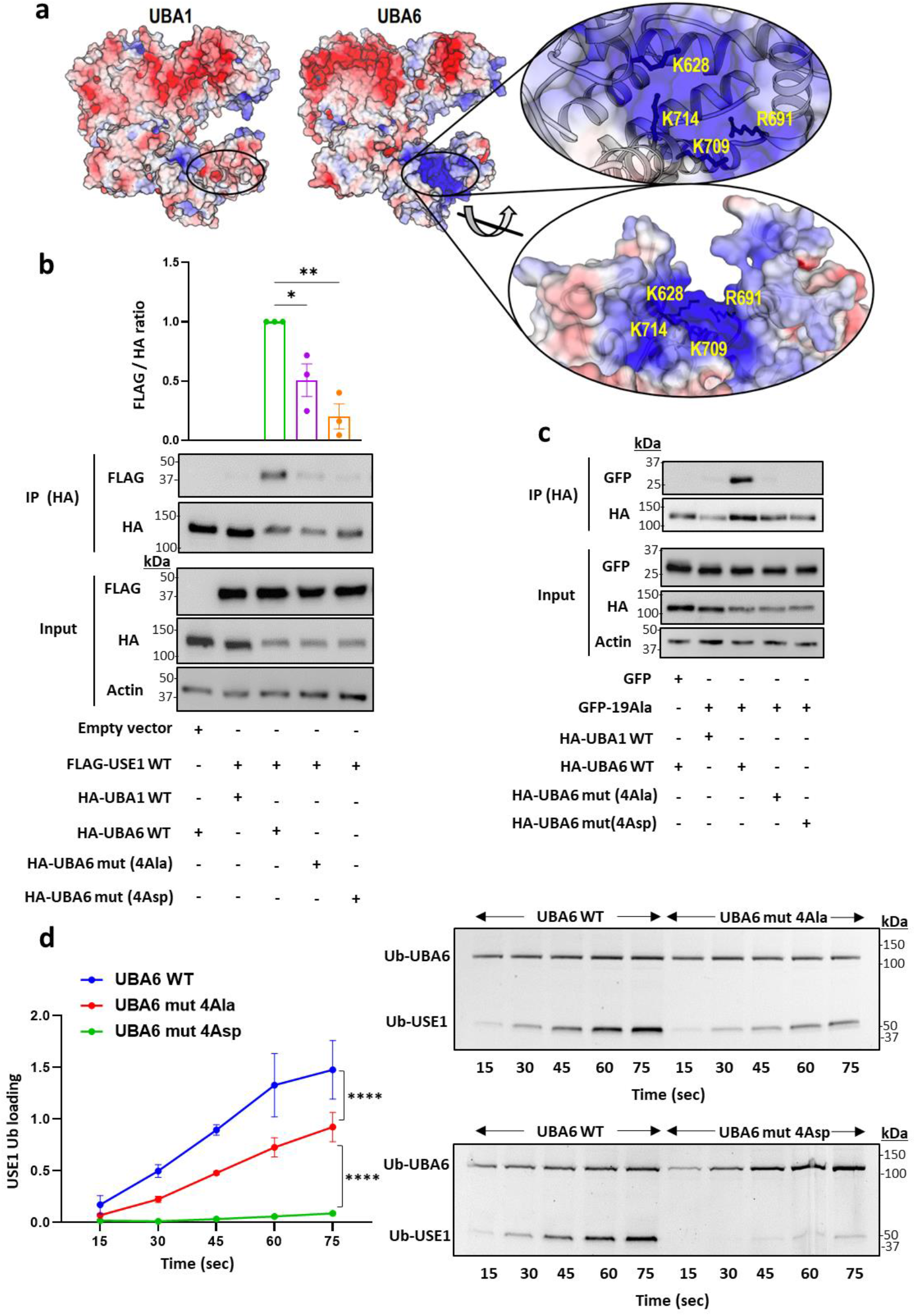
UBA6 interacts with polyalanine stretches. **a**, Electrostatic surface representation of the UBA6 structural model in comparison to UBA1. The location of key Arg and Lys residues forming the positively charged patch in UBA6 is presented. **b-c**, HA-tagged constructs of WT UBA1, WT UBA6, and UBA6 mutants with Ala or Asp substitution mutations in Lys628, Arg691, Lys709, and Lys714 (UBA6 mut 4Ala or UBA6 mut 4Asp) were co-transfected with FLAG-USE1 (**b**) or empty GFP and GFP-polyAla (19Ala) (**c**) into HEK293T cells. Cell lysates were immunoprecipitated with anti-HA antibodies and the immunocomplexes were analyzed with anti-FLAG, anti-HA, and anti-GFP antibodies. Results are mean ± s.e.m. One-way ANOVA Dunnett’s test, n=3. **d**, Representative gel of time-dependent *in vitro* ubiquitin loading of USE1 by WT UBA6, UBA6 mut 4Ala, and UBA6 mut 4Asp in the presence of fluorescein-labelled ubiquitin. Imaging of fluorescent ubiquitin conjugates was carried out with a laser scanner at 488 nm. Quantification of USE1 ubiquitin loading is presented as mean ± s.e.m. Two-way ANOVA Tukey’s test, n=3. \**P* < 0.05, \*\**P* < 0.01, **** *P* < 0.0001.

### Disease-causing proteins with polyalanine expansions interact with UBA6

Since UBA6 is mainly localized in the cytoplasm in HEK293T cells (**Extended Data Fig. 3a**), we next expressed isolated polyalanine stretches with or without a nuclear localization sequence (NLS) and monitored their binding to UBA6. As expected, polyalanine stretches bound UBA6 effectively when expressed without NLS (**Fig. 3a**), and reduced the ubiquitin loading of USE1 (**Extended Data Fig. 4a**). However, in contrast, isolated polyalanine stretches with NLS did not bind UBA6 and had no apparent effect on USE1 loading (**Fig. 3a**, **Extended Data Fig. 4a**). To explore pathological UBA6 interactions, we expressed various disease-causing proteins with polyalanine expansion mutations in cells (**Extended Data Fig. 3 b-e**) and compared their interactions with UBA6 to those of the wild type protein (**Fig. 3** **b-e**). This includes PHOX2B (WT, +13 Ala, **Fig. 3b**), RUNX2 (WT, +6 Ala, +12 Ala, **Fig. 3c**), HOXD13 (WT, +10 Ala, **Fig. 3d**), and PABPN1 (WT, +7 Ala, **Fig. 3e**).

**Figure 3.**
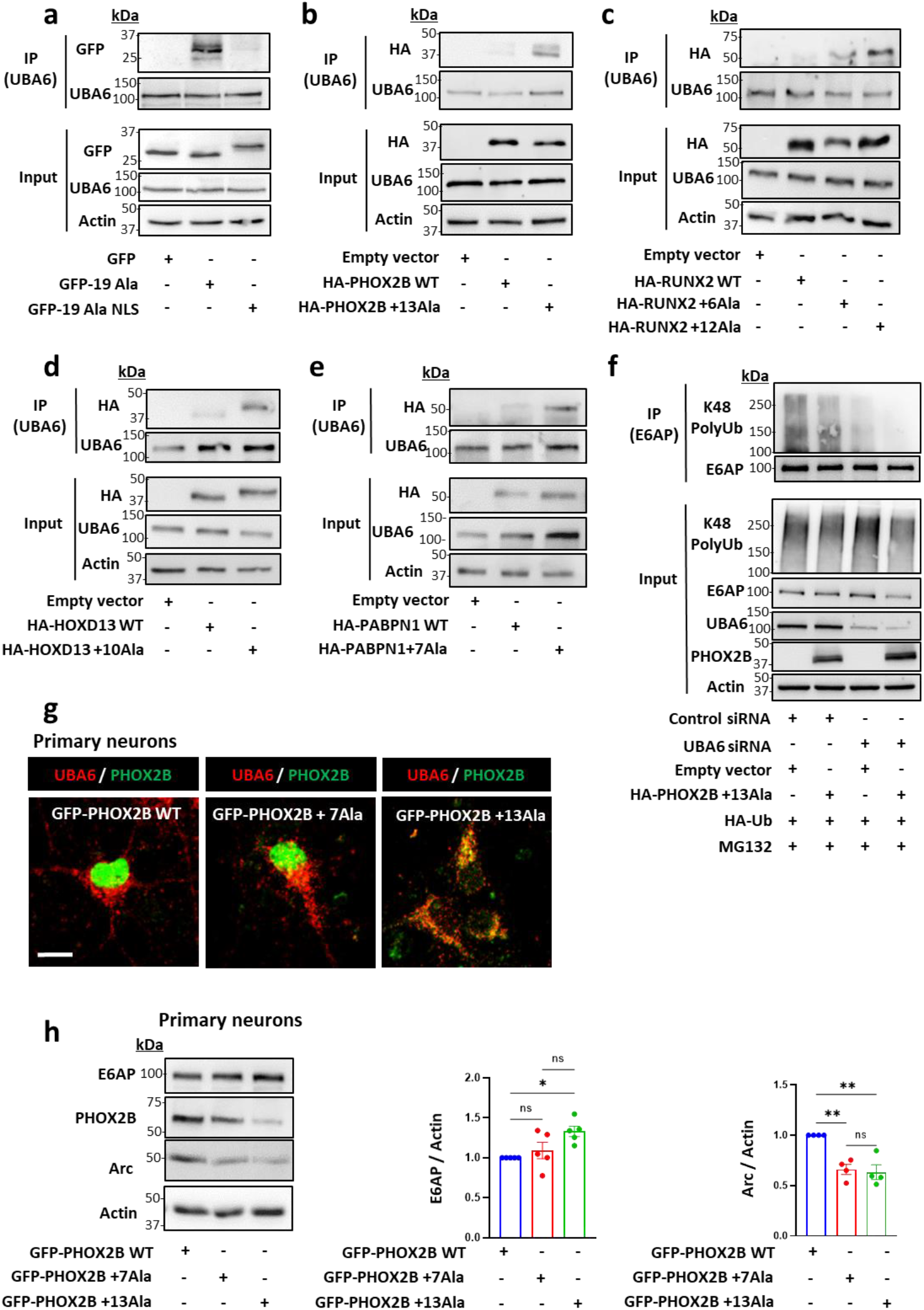
Polyalanine-expanded disease proteins interact with UBA6 and inhibit E6AP degradation. **a-e**, HEK293T cells were transfected with the indicated constructs and immunoprecipitated for endogenous UBA6. **a**, Empty GFP, and GFP-polyAla with or without a nuclear localization sequence (NLS). **b**, WT and mutant PHOX2B (+13Ala). **c**, WT and mutant RUNX2 (+6Ala and +12Ala). **d**, WT and mutant HOXD13 (+10Ala). **e**, WT and mutant PABPN1 (+7 Ala). **f**, Control and UBA6-depleted HEK293T cells were transfected with HA-Ub, mutant PHOX2B or empty vector and incubated for the last 6 h with the proteasome inhibitor MG132 (10 μM). Endogenous E6AP was immunoprecipitated from cell lysates for ubiquitination analysis. **g-h,** Mouse primary cortical neurons were transduced with lentiviral vectors expressing GFP-tagged WT PHOX2B or mutant PHOX2B (+7 Ala and +13Ala). **g,** Association of endogenous UBA6 with GFP-PHOX2B. Scale bar 10 μm. Quantification is shown in **Extended Data Fig. 5c**. **h**, Analysis of E6AP and Arc levels in the WT and mutant PHOX2B-expressing neurons (n=5 and n=4, respectively). Results are mean ± s.e.m normalized to WT PHOX2B. One-way ANOVA Tukey’s test, ns non-significant, \**P* < 0.05, \*\**P* < 0.01.

These pathological interactions are measurable because although UBA6 is mainly localized to the cytoplasm and, as expected, wild type PHOX2B, RUNX2, HOXD13, and PABPN1 are primarily localized in the nucleus, all the mutant proteins with polyalanine expansions partially mislocalized to the cytoplasm and could then interact with endogenous UBA6 (**Extended Data Fig. 3 b-e**, **Fig. 3 b-e**).

### Inhibition of UBA6-depndent E6AP degradation by polyalanine expansions

E6AP/UBE3A is a highly potent E3 ubiquitin ligase whose regulation is critical for normal development of the nervous system. Indeed, decreased activity of the ligase results in Angelman syndrome while increased activity causes autism spectrum disorders ^14, 24, 25^. Harper and co-workers identified a unique regulatory cascade in which UBA6-USE1 ubiquitinates E6AP for degradation in the proteasome ^13^. Accordingly, our results indicate that depletion of UBA6, as well as expression of mutant PHOX2B (+13 Ala), increase the levels of E6AP (**Extended Data Fig. 4b**). This correlates with a decrease in Lys48-linked polyubiquitination of E6AP (**Fig. 3f**), suggesting that proteasome-mediated degradation of E6AP is inhibited by mutant PHOX2B. Moreover, E6AP levels were increased in the polyalanine-expressing cells, which was reversed by UBA6 overexpression (**Extended Data Fig. 4c**), compatible with a model in which the polyalanine stretch impairs the regulation of E6AP levels by competing with USE1 on UBA6.

In order to test our model in neuronal cells, we used mouse primary cortical neurons. Similarly, UBA6 was predominantly detected in the cell body and in neurites, with only a small fraction localized to the nucleus (**Extended Data Fig. 5a**). Transduction of the neurons with lentiviruses expressing GFP-tagged wild type or mutant PHOX2B (+13 Ala) resulted in a significant difference in subcellular localization and expression pattern. While most of the wild type PHOX2B could be detected in the nucleus, mutant PHOX2B (+13 Ala) was also found in the cell body and in neurites, where the GFP fluorescence imaging revealed both non-aggregated and aggregated patterns (**Extended Data Fig. 5b**). Neurons expressing mutant PHOX2B showed increased colocalization with UBA6, which correlated with increasing lengths of the polyalanine stretch (**Fig. 3g, Extended Data Fig. 5c**). Moreover, expression of mutant PHOX2B with +7 Ala and +13 Ala increased the levels of E6AP in the primary neurons (**Fig. 3h**). The synaptic activity-regulated cytoskeleton-associated protein (Arc) is negatively regulated by E6AP ^26, 27^, and accordingly, the increased levels of E6AP due to ectopic expression of the PHOX2B mutants with +7 Ala and +13 Ala was accompanied with a decrease in Arc levels in primary neurons (**Fig. 3h**, **Extended Data Fig. 5d**).

### UBA6 dysregulation in neurons of patients with polyalanine expansions

In order to investigate the mechanisms underlying human autonomic neuron vulnerabilities, we sought to assess the associations between endogenous polyalanine expansions and UBA6. Endogenous PHOX2B is expressed in specific neuronal populations of the autonomic nervous system ^28, 29^. In humans, polyalanine expansion mutations within the *PHOX2B* gene cause CCHS, a rare and life-threatening condition with autonomic nervous system dysfunction ^30, 31^. We have previously described the generation of two induced pluripotent stem cells (iPSCs) from identical twins carrying a heterozygous PHOX2B +5 Ala expansion (101iCCHS 20/25 and 102iCCHS 20/25) ^32^. Here, we obtained skin punch biopsies from an additional CCHS patient bearing a heterozygous +7 Ala expansion (104iCCHS 20/27), as well as from their sex-matched healthy family relatives, which serve as controls (103iCTR 20/20 and 105iCTR 20/20) (**Extended Data Fig. 6**). These patients suffer from central sleep apnea with a more severe peripheral autonomic presentation in 104iCCHS 20/27, including Hirschsprung disease. Patient-derived fibroblasts were reprogrammed using non-integrating episomal plasmids as previously described ^32^. All generated iPSC lines expressed the pluripotency markers NANOG, SOX2, OCT3/4 TRA-1-60, and SSEA4 (**Extended Data Fig. 6 a-j**), and spontaneously differentiated into the three germ layers (**Extended Data Fig. 6 l-n**). All lines showed a normal karyotype, and genetic analysis confirmed the presence of a heterozygous expansion of seven alanine residues resulting with a 27 polyalanine stretch in PHOX2B (**Extended Data Fig. 6 k, o**). In order to obtain endogenous PHOX2B-expressing cells, we differentiated the iPSCs to neural crest progenitor cells that were further differentiated into sympathetic neuroblast crest cells, and finally to peripheral autonomic neurons (**Fig. 4a**). Characterization of the human autonomic neurons revealed expression of PHOX2B, the pan-neuronal marker βIII-tubulin, the catecholaminergic marker tyrosine hydroxylase (TH), the peripheral neuronal marker peripherin, and atonal BHLH transcription factor 1 (ATOH1), which is associated with breathing and digestion (**Fig. 4a**).

**Figure 4.**
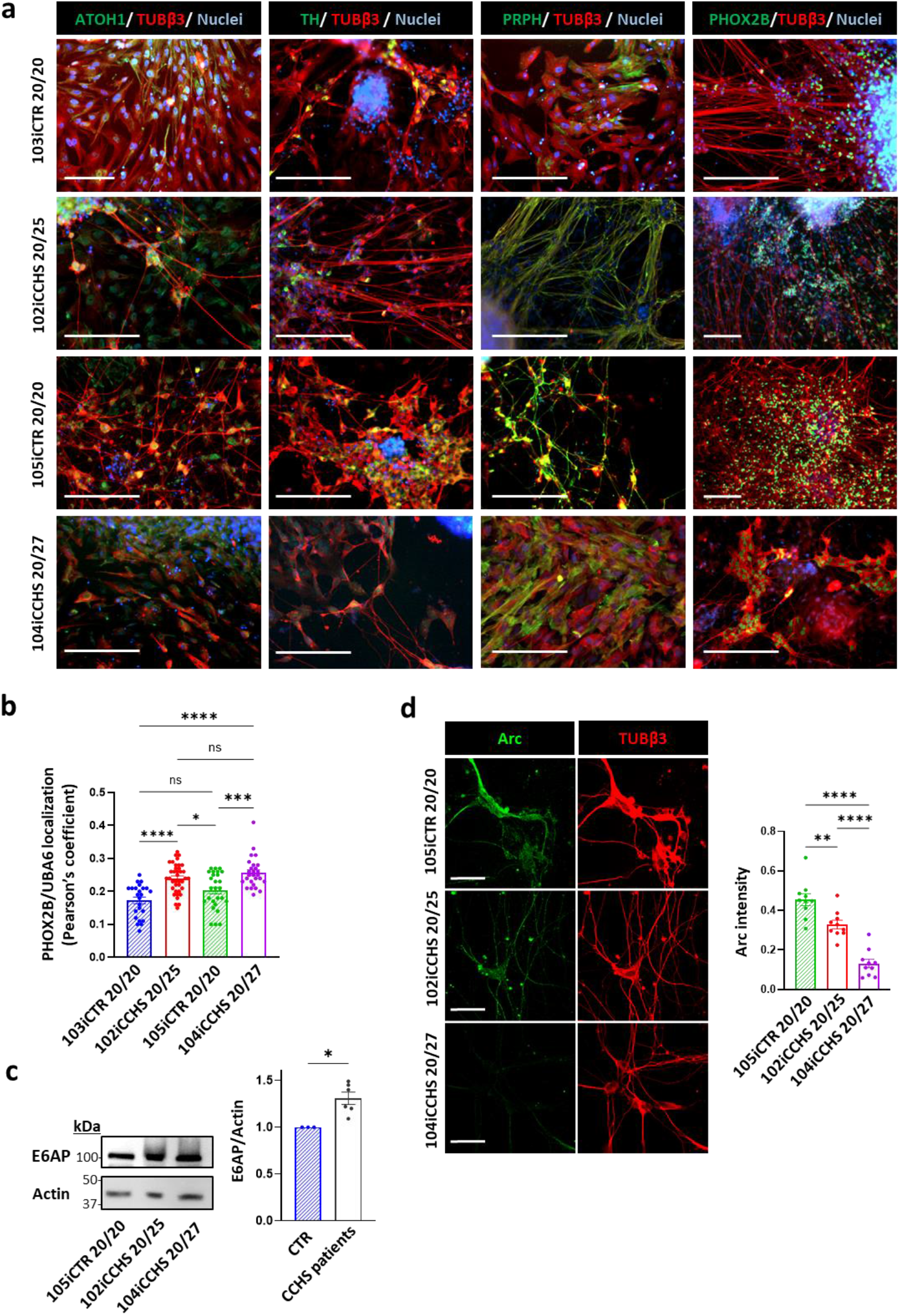
iPSC-derived autonomic neurons from CCHS patients showing UBA6-dependent alterations in E6AP and Arc levels. **a,** Characterization of iPSC-derived autonomic neurons at day 31 of differentiation from healthy controls and CCHS patients. Immunocytochemistry of PHOX2B, βIII-tubulin (TUBβ3), tyrosine hydroxylase (TH) peripherin (PRPH), and atonal BHLH Transcription Factor 1 (ATOH1). Scale bar 200 μm. **b**, Quantification of the association of endogenous UBA6 with endogenous PHOX2B (Pearson’s coefficient) in autonomic neurons from control and CCHS patients. Results are the average values from neurons in different imaged fields. Total number of neurons analyzed was 160 for 103iCTR 20/20, 350 for 102iCCHS 20/25, 380 for 105iCTR 20/20 and 550 for 104iCCHS 20/27. One-way ANOVA Tukey’s test. Images are shown in **Extended Data Fig. 7c**. **c,** Analysis of E6AP levels in control (105iCTR 20/20) and CCHS patients (102iCCHS 20/25 and 104iCCHS 20/27). Results are mean ± s.e.m normalized to control from three independent differentiation days. Unpaired 2-tailed t-test. **d,** Immunostaining of Arc and TUBβ3 in autonomic neurons from control and CCHS patients. Scale bar 50μm. For quantification, Arc intensity was normalized to TUBβ3. Results are mean ± s.e.m from 40 analyzed neurons. One-way ANOVA Tukey’s test, ns non-significant, \**P* < 0.05, \*\**P* < 0.01, \*\*\**P* < 0.001, \*\*\*\**P* < 0.0001.

Analysis of the PHOX2B localization showed a decrease in the nuclear fraction of PHOX2B in the mutant 20/25 and 20/27 neurons, as compared to healthy control neurons (**Extended Data Fig. 7 a,b**). The cytoplasmic PHOX2B was mainly perinuclear without visible aggregates (**Extended Data Fig. 7c**). In addition, there was an increased association between UBA6 and PHOX2B in the CCHS patient neurons (**Fig. 4b**). Some CCHS neurons exhibited severe PHOX2B cytoplasmic mislocalization, with apoptotic nuclear morphology (1.9% and 3.9% of the 20/25 and the 20/27 mutations, respectively) (**Extended Data Fig. 7c**). In accordance with the mouse data, E6AP levels were higher in the patient neurons (**Fig. 4c**). Moreover, Arc levels were lower in the CCHS neurons with the 20/25 mutation than in control cells and the levels were further reduced in CCHS neurons with the 20/27 mutation (**Fig. 4d**).

## Discussion

Expansion mutations in stretches of repetitive DNA sequences that encode poly amino acids such as polyglutamine or polyalanine can cause various diseases. Whereas much progress has been made on the molecular mechanisms of polyglutamine diseases ^9, 33, 34^, the consequences of polyalanine expansions remain more of an enigma. Here, we demonstrate how expanded polyalanine stretches can compete with the normal function of a shorter stretch to disrupt the specific interaction between the E2 ubiquitin-conjugating enzyme USE1 and the E1 ubiquitin-activating enzyme, UBA6. Our results indicate that UBA6 interacts with polyalanine stretches, and that this interaction contributes to UBA6 recognition of USE1, but can be altered by different polyalanine disease-causing proteins.

The correlation between polyalanine expansion mutations and a possible harmful gain-of-function is supported by the observation that the respiratory abnormalities of *phox2b^+/-^*mouse mutants are transient and milder than those exhibited by *phox2b^+/27Ala^*mouse mutants or those of CCHS patients ^35, 36^. These genetic studies suggest the possibility of additional disease contributing mechanisms besides partial loss of PHOX2B function in the nucleus and alterations in transcriptional programs by polyalanine expansion mutations ^37^. UBA6 plays an important role in mouse embryonic development, neuronal function, and survival ^13, 38^. Thus, the inhibition of UBA6 by sequestering the enzyme into cytoplasmic mislocalized polyalanine expansions may represent a deleterious mechanism for such mutations. The correlation between disease severity and the length of the polyalanine expansion may be related to the greater tendency of mutants with longer polyalanine stretches to mislocalize to the cytoplasm and interact there with UBA6. This tendency of polyalanine-expanded proteins to mislocalize to the cytoplasm has been previously detected in overexpression systems ^10^, but the relevance to human disease was questioned. Our results provide evidence that cytoplasmic mislocalization of endogenous PHOX2B can occur in the neurons affected in CCHS, which may contribute to the neuronal dysfunction seen in this disease.

In summary, our study provides mechanistic insights into the long-standing question of the functional role of polyalanine stretches and demonstrates that they are relevant to the ubiquitin system and its dysregulation in disease states.

## Methods

### Ethics statement

All mouse experiments were reviewed and approved by the Institutional Animal Care and Use Committee of Tel Aviv University. All cell lines and protocols related to human stem cell research in the present study were used in accordance with guidelines approved by the Institutional Review Board (Sheba Medical Center and Tel Aviv University). Informed consent was obtained from all donors.

### Antibodies

The following antibodies were used in this study: mouse anti-HA (Biolegend 901501); rabbit anti-HA (Cell signaling 3724); rabbit anti-Actin (SIGMA A2066); mouse anti-E6-AP (Santa Cruz 166689); mouse anti-FLAG (SIGMA F1804); rabbit anti-UBA6 (Cell signaling 13386); rabbit anti-K48 polyUb (Cell signaling 8081S); rabbit anti-UBE2Z (USE1, (Abcam 254700); rabbit anti-GFP (Abcam 6556), mouse anti-PHOX2B (Santa Cruz 376997); mouse anti-mCherry (Abcam 125096); mouse anti-Tau (Santa Cruz 58860); rabbit anti-MAP2a (synaptic systems 188003); mouse anti-Arc (BD Transduction Laboratories 612602); rabbit anti-Arc (Abcam 183183); Alexa Fluor 555 (Abcam 150114) conjugated goat anti-mouse secondary antibody, Alexa Fluor^®^ 555 (Abcam 150078) conjugated goat anti-rabbit secondary antibody, goat anti-mouse (Abcam 6789) and goat anti-rabbit (Abcam 6721); horseradish peroxidase (HRP)-conjugated secondary antibodies, mouse anti-MATH-1 (Santa Cruz 136173), mouse anti-PRPH (Santa Cruz 377093); mouse anti-TH (Santa Cruz 25269); mouse anti-SOX10 (Abcam 155279); rabbit anti-β-Tubulin III (SIGMA T2200); mouse anti-OCT3/4 (Santa Cruz 5279); mouse anti-TRA 1-60 (R & D MAB4770); mouse anti-SSEA-4 (Santa Cruz 21740); mouse anti-IgG3 Isotype (Bio-legend 330405); rabbit anti-SOX2 (Abcam 97959); rabbit anti-NANOG (Abcam 21624); rabbit anti-68kDa Neurofilament (Abcam 52989); rabbit anti-α-Fetoprotein (ScyTek A00058); rabbit anti-α-smooth muscle actin (Abcam 32575); Alexa Fluor 488-conjugated anti-mouse secondary antibody (Jackson Immuonoresearch 715545150); Alexa Fluor 488-conjugated anti-mouse secondary antibody (Thermo Fisher Scientific, A-11034); Alexa Fluor 488-conjugated anti-rabbit secondary antibody (Molecular Probes, A-11029); Alexa Fluor 594-conjugated anti-rabbit secondary antibody (Jackson Immuonoresearch 711585152); Alexa Fluor 568-conjugated anti-rabbit secondary antibody (Abcam 175471); and Alexa Fluor 647-conjugated anti-rabbit secondary antibody (Abcam 150079).

### Cloning and constructs

E1 constructs for mammalian cell expression, namely pcDNA3.1-HA-tagged UBA1 wild type and pcDNA3.1-HA-tagged UBA6 (UBE1L2) wild type, were kindly provided by Dr. Marcus Groettrup. pcDNA3.1-HA-UBA6 mut (4Ala) and pcDNA3.1-HA-UBA6 mut (4Asp) where Lysine 628, 709,714 and Arginine 691 were changed to amino acids Alanine and Aspartate respectively, were constructed by Gibson assembly using the DNA gene blocks. For *E. coli* expression as a His6-tagged proteins, the wild type and mutant UBA1 and UBA6 genes where subcloned into modified and improved vector pET28a ^39^.

E2 USE1 (UBE2Z) expressing plasmids pcDNA3.1-His6-3xFlag-USE1 was kindly provided by Dr. Annette Aichem. For expression in mammalian cells, the putative Kozak sequence was first added to the USE1 gene, which was subcloned into pEGFP-N1 based plasmid resulting in plasmid pCMV-His6-3xFlag-USE1 new Kozak used in this study. The ΔPolyAla region mutant, where amino acids 47-56 of the USE1 gene were deleted, the 2A>2R mutant where Alanine 49 and Alanine 52 were changed to Arginine respectively, the ΔLB mutant where amino acids 194-197 comprising a Loop B ^16^ were deleted, and the C188A mutant were constructed by site-directed mutagenesis and Gibson assembly applying Q5® Site-Directed Mutagenesis Kit and NEBuilder® HiFi DNA Assembly respectively. For *E.coli* expression as a His6-tagged proteins, the USE1 wild type and mutant genes were subcloned into modified plasmid pET30-His6-N-AviTag.

His6-Ub plasmid for expression in *E. coli* of the His-tagged ubiquitin gene was described previously ^40^. The HA-ubiquitin plasmid for mammalian expression was a gift from Edward Yeh (Addgene plasmid # 18712 ^41^).

pEGFP-19 Alanines and pEGFP-19 Alanines with nuclear localization sequence (NLS) constructs for expression in mammalian cells as well as HA-bovine PABPN1 wild type and HA-bovine PABPN1 mut +7 Ala mutant constructs were a gift of Dr. David Rubinsztein. First, a delta polyalanine region mutant was constructed by Inverse PCR and then the PABPN1 gene was transferred into pEGFP-N1 based vector. In order to construct human PABPN1 wild type and mutant genes, a site directed mutagenesis was applied to change amino acids Asparagine 95 to Serine and Serine 102 to Proline on the delta Polyalanine region construct. Humanized PABPN1 wild type and +7 Ala mutant plasmids were constructed by adding polyalanine regions applying the DNA gene blocks and Gibson assembly.

Human RUNX2 carrying plasmids pcDNA3.2/GW/D-TOPO RUNX2 wild type, pcDNA 6.2/C-EmGFP-DEST-RUNX2 mut (+6 Ala) and pcDNA 6.2/C-EmGFP-DEST-RUNX2 mut (+12 Ala) were kindly provided by Dr. Yoshihito Tokita. In this study, in all aforementioned plasmids, the EmGFP gene was deleted and HA-tag was introduced onto the C-terminus of the RUNX2 genes.

Mouse HOXD13 bearing plasmids pcDNA3.1-Hoxd13 wild type and pcDNA3.1-Hoxd13 mut +10Ala were a gift of Dr. Denes Hnisz. First, the Valine259 to Glutamate mutation was corrected in both constructs and then the wild type and the mutant Hoxd13 genes were subcloned into pEGFP-N1 derived vector whereas adding a putative Kozak sequence and the C-terminal HA-tag.

PHOX2B carrying plasmids pcDNA3.0-HA-PHOX2B wild type and pcDNA3.0-HA-PHOX2B mut (+13 Ala) were kindly provided Dr. Diego Fornasari. To express the PHOX2B genes alone and as fusion with EGFP in neuronal cells under control of the human Synapsin I promoter, the PHOX2B genes were subcloned into lentiviral pLL3.7 vector bearing the Synapsin I promoter. Plasmids bearing the HA-PHOX2B mut (+7Ala) gene were constructed applying Gibson assembly and resulted in plasmids pcDNA3.0-HA-PHOX2B mut (+7 Ala), pLL3.7-hSyn-HA-PHOX2B mut (+7 Ala) and pLL3.7-hSyn-HA-PHOX2B mut (+7 Ala)-EGFP.

The integrity of every construct used in this study was verified by the sequencing analysis and the detailed explanations of cloning procedures will be provided upon request.

### Cell lines and transfection

Cell lines used in this study include human embryonic kidney cells, HEK293T (ATCC CRL-1573) and HEK293FT (Invitrogen, R70007). The cells were authenticated by STR profiling and were routinely tested for mycoplasma contamination. The cells were grown in Dulbecco’s modified Eagle’s medium (01-052-1A, Biological Industries) supplemented with 10% heat-inactivated fetal bovine serum (04-007-1A, Biological Industries), 10000 units/mL penicillin, and 10 mg/mL streptomycin (03-031-1B Biological Industries) and 2 mM L-glutamine (G7513, SIGMA) at 37 °C with 5% CO_2_. Cells were seeded and cultured for approximately 24 h until they grew to 50– 60% confluence before transfection. Transient transfection of indicated plasmids, was accomplished using TransIT-LT1(Mirus, MIR 2300) according to the manufacturer’s protocol. Vector and Mirus were mixed in a reduced serum medium (Opti-MEM^®^ 10001865, Gibco) and incubated at room temperature for up to 30 min before being dripped gently onto the cell culture and incubated for 24 to 48 h. Transfection efficiency was confirmed by western blot analysis. In RNA interference experiments, cells were transfected 24 h after seeding with 50-100 nM *SMART*pool siRNAs (Dharmacon) for gene silencing and Lipofectamine 2000 (1000186, Invitrogen), with two rounds of knockdown for 5 days, according to the manufacturer’s instructions (Invitrogen). For this purpose, the siRNAs and Lipofectamine were diluted separately in reduced serum medium Opti-MEM® (10001865, Gibco), then mixed for 15 min at room temperature and dripped gently onto the cell culture, which was then incubated at 37 °C for 4-6 h, before restoration of full medium. The following oligonucleotides (ON-TARGETplus SMARTpool, Dharmacon L-006403-01-0005) were used for UBA6 depletion:

siRNA J-006403-09, UBA6: Target Sequence: GUGUAGAAUUAGCAAGAUU

siRNA J-006403-10, UBA6: Target Sequence: GCAUAGCUGUCCAAGUUAA

siRNA J-006403-11, UBA6: Target Sequence: CAGUGUUGUAGGAGCAAUA

siRNA J-006403-12, UBA6: Target Sequence: GGAAUUUGGUCAGGUUAU

### Isolation and culture of mouse primary cortical neurons

Primary cortical neurons were isolated from wild-type C57BL/6J mouse embryos at E17. Brains were harvested and placed in ice-cold HBSS under a dissection microscope. Cerebral cortices were dissected and incubated in HBSS. After mechanical dissociation using sterile micropipette tips, dissociated neurons were resuspended and cultured at 37°C in a humidified incubator with 5% CO_2_ and 95% O_2_ in poly-D-lysine coated 6-well plates in neurobasal media (12349015, Gibco) supplemented with 1% GlutaMAX™ Supplement (35050-061, Thermofisher), 1% Sodium pyruvate (11360039, Gibco), 2% B27 supplement (17504044, Gibco) and 1% Penicillin-Streptomycin (03-031-1B, Biological Industries). One-half of the culture media was changed every three days until treatment. Differentiated cortical neurons were infected with indicated lentiviral vectors after 5 days in culture.

### Lentivirus production and infection

Third generation lentiviral vectors pLL3.7 (Addgene, #11795) that express shRNA under the mouse U6 promoter and CMV-EGFP or hSyn-EGFP reporter cassettes were obtained from the TAU Viral Core facility. Helper plasmids pMDLg/pRRE, pRSV-Rev, and pCMV-VSVG that carry HIV regulatory protein genes as well as the pseudotyped envelope protein gene from vesicular stomatitis virus envelope G (VSV G), were obtained from the same facility.

Briefly, HEK-293FT packaging cells growing in 15 cm dishes were transfected with a mix of 7.8 μg helper vector pMDLg/pRRE, 3 μg helper vector pRSV-Rev, 4.2 µg envelope vector pCMV-VSVG, and 12 µg target vector pLL3.7-hSyn-PHOX2B-EGFP carrying the wild type or mutant PHOX2B gene. Polyethylenimine (PEI), (SIGMA 408727) was used as a transfection reagent. At 16 hours’ post-transfection, the culture media was removed and replaced with fresh high-serum medium, which was harvested 48 h later and filtered through Amicon Ultra -15 (UFC910024) vials at 1500g for 30 minutes to obtain concentrated and purified lentiviruses for transduction. On the day of transduction, half the culture media was removed from primary cultured cortical neurons and was stored for later use (conditioned media), and then viral particles were added and incubated for 15 hours. At the end of this time, the culture medium was replaced by a 1:1 mix of fresh and the conditioned medium that was collected previously. The neurons were then incubated for five more days before being harvested for either western blot or immunostaining.

### Generation of patient-specific fibroblasts

Skin biopsies were collected from a 2-year-old female CCHS patient who carries a 20/27 Ala PHOX2B genotype as well as from her healthy sister and from the healthy father of two previously reported identical 4-year-old twin males with CCHS who carry 20/25 Ala stretches ^32^. The biopsies were dissected and cultured for two weeks in 6-well plates under a coverslip in DMEM (Biological industries 011701A) with 20% FBS and with half the media replaced every other day.

### Reprogramming of iPSCs

One million fibroblast cells were harvested using TrypLE Express (Gibco, 12604021) and electroporated with non-integrating episomal vectors using a Neon transfection system (Invitrogen, kit MPK10096). The cells were then plated on mouse embryonic fibroblast (MEF)-coated plates and cultured in DMEM with 15% FBS, 5 ng/ml basic fibroblast growth factor (bFGF, Peprotech 10018B) and 5 μM ROCK inhibitor (Enzo, ALX270333). After two days, the medium was replaced with NutriStem (BI) supplemented with 5 ng/ml bFGF, with fresh medium added every other day. On day 22, six colonies were transferred to new MEF-coated plates and cultured in NutriStem with 5 ng/ml bFGF. Three colonies were selected and manually transferred to matrigel (Corning)-coated plates and cultured in NutriStem, which was replaced daily and with weekly passage.

### iPSC characterization

iPSCs were assessed for the expression of the pluripotency markers NANOG, SOX2, OCT3/4, TRA 1-60, and SSEA by immunocytochemistry and FACS analysis. The differentiation potential was assessed by harvesting the iPSCs at confluency using TrypLE and resuspending the cells in NutriStem supplemented with 10 ng/ml bFGF and 7 μM ROCK inhibitor (Enzo, ALX270333). Embryoid bodies (EBs) spontaneously formed after 2 days, at which time, the medium was replaced with EB medium (DMEM with 15% FBS, 1% Non-Essential Amino Acids and 0.1 mM β-mercaptoethanol, Gibco 31350010). After 4–7 days, the EBs were plated on 0.1% gelatin-coated plates and cultured for 21 days with EB medium replacement twice weekly. On day 21, the cells were fixed and stained for heavy chain neurofilament, α-SMA, or α-fetoprotein as ectodermal, mesodermal and endodermal markers, respectively. G-banding karyotype analysis was used to exclude any chromosomal abnormalities that may have occurred during the reprogramming process. Briefly, iPSCs were supplemented with 100 ng/ml colcemid (Biological Industries 120041D), incubated for 60 min, and harvested in Versene solution (Gibco 15040033). Cells were fixed in 1:3 glacial acetic acid:methanol (Biolabs-chemicals) solution and the G-banding karyotype was determined. Finally, all lines were tested for mycoplasma contamination using the Hy-mycoplasma PCR kit (Hylabs, KI5034I).

### Differentiation of iPSCs into autonomic neurons

The protocol was as described previously ^42^. The iPSCs were cultured in Nutristem to a confluence of 1 million cells/well. On day 0, a single cell suspension was prepared using Versene solution (Gibco, 15040033), and then the cells were quickly re-aggregated in T25 flasks coated with Poly-Hema (2-hydroxyethyl methacrylate) at a concentration of 350,000 cells/ml and were cultured in neuromesodermal progenitor cell induction medium (NMP media) containing Essential 6 media (Gibco, A1516401), 1.5 mM CHIR (Tocris biotech, 4423), 10 μM SB (Tocris biotech, 431542), Penicillin-Streptomycin-Amphotericin B solution (PSA, Biological Industries, 03-033-1B), and 10 μM ROCK inhibitor (ROCKi Enzo, ALX270333).

On day 1, half the media were replaced with medium without ROCKi, and on day 3, the media were replaced with sympathetic neural crest induction medium (NCi medium) containing Essential 6 media, 1.5 mM CHIR, 20 ng/ml bFGF (Peprotech 10018B), 50 ng/ml BMP4 (Prospec, Cyt-1093), 100 nM all trans retinoic acid (SIGMA, R2625), and PSA. On day 10, the culture was dissociated into single cells using Accutase (Gibco, A1110501) for 4 min at 37°C, and cultured in sympathetic neuroblast induction and propagation (NCC) medium, containing neurobasal media (Gibco 21103049), 20 ng/ml bFGF, 50 ng/ml BMP4, 20 ng/ml EGF, 2 μg/ml heparin, B27 (Gibco 12587010), N2 (Gibco 17502048), GlutaMAX (Gibco 35050038) and PSA. On day 11, half the NCC media was replaced by medium without ROCKi and half the media were subsequently replaced every other day. On day 17, the medium was replaced with sympathetic neuronal maturation medium (NMM medium) containing neurobasal media, B27, N2, 10 ng/ml GDNF (Peprotech 45010), 10 ng/ml BDNF (Peprotech 45002), 10 ng/ml NGF (R & D 256-GF), GlutaMAX, and PSA. One third of the media were subsequently replaced every other day until day 31.

### Immunocytochemistry for iPSCs-derived autonomic neurons

In order to prepare for the culture, coverslips were incubated in poly-L-ornithine solution (SIGMA, P3655) overnight at 37°C followed by 3 washes with cell culture grade water and drying for 15 min. On seeding day, the coverslips were incubated in laminin (Merk, L2020) d for 1 h at 37°C before being washed with PBS. The neurospheres were harvested manually and seeded on the coverslips. After 5-4 days, the cells on the coverslips were washed with Dulbecco’s Phosphate-Buffered Saline (DPBS, Biological Industries, 020231A) and fixed in 4% paraformaldehyde at room temperature for 15 min. The cells then were washed twice in DPBS, and blocked with DPBS containing 0.1% Triton X-100 (SIGMA, T8532) and 1% bovine serum albumin (blocking solution, SIGMA, A7906100G), for 1 h at RT. Primary antibodies were added to the blocking solution and incubated overnight at 4°C. Following three washes with blocking solution, the cells were incubated with fluorescent secondary antibodies for 2 h at room temperature. DRAQ5 was used to stain the cell nuclei.

### Flow cytometry

iPSCs were harvested and dissociated into single cells by incubation with TrypLE for 2 min at 37°C. For the detection of intracellular markers, samples were incubated in fixation solution (Invitrogen, 00522356, 00512343) for 40 min at room temperature followed by washings with permeabilization solution (Invitrogen, 00833356). For surface markers, the cells were washed once with 3% FBS in DPBS. In both cases, the samples were incubated with the appropriate primary antibodies for 2 h at room temperature, then were washed twice and incubated for 1 h with the relevant secondary antibody followed by three more washes. Analysis was performed using a NovoCyte flow cytometer (ACEA). The first gating was SSC-H/FSC-H, and the entire cell population was selected (without cell debris) followed by FSC-H/FSC-A gating and single cell selection. The final analysis is count (%)/FITC-H.

### Western blotting assay

Cells were washed with PBS and harvested in Laemmli buffer containing 5% beta-mercaptoethanol. For the ubiquitin loading assays, the cells were lysed on ice in lysis buffer (20 mM Tris-HCl, pH 6.8, 137 mM NaCl, 1 mM EGTA, 1% Triton x100, 10% glycerol, and a protease inhibitors cocktail), centrifuged to discard the cell pellet and then the supernatant was added to Laemmli buffer at a ratio of 1:1 without using beta-mercaptoethanol. Protein samples were boiled for 5 min at 95°C, separated by SDS–PAGE, transferred onto PVDF membranes, subjected to western blot analysis, and visualized using the ECL enhanced chemiluminescence reagent (CYANAGEN). Protein levels in each sample were evaluated by normalization to the housekeeping β-actin. The bands were quantified using ImageJ software.

### Immunoprecipitation and ubiquitination assays

Cells were lysed on ice in IP buffer (20 mM Tris-HCl, pH 7.2, 150 mM NaCl, 2 mM MgCl_2_, 0.5% NP-40), supplemented with a protease inhibitors cocktail before use. For ubiquitination experiments, cells were treated with a proteasome inhibitor MG132 (10 μM) during the last 6 hours before lysis with the IP buffer supplemented with 1 mM PMSF and 10 mM iodoacetamide. Whole-cell lysates obtained by centrifugation were incubated with 2-5 μg of antibody overnight at 4°C, followed by 2 h incubation with Protein A-Sepharose CL-4B (GE-Healthcare, 17-0780-01). The immunocomplexes were then washed three times with IP buffer, and boiled at 95°C for 5 minutes in Laemmli sample buffer, before being separated by SDS–PAGE for western blotting assays. For experiments to examine the binding of isolated polyalanine stretches, a pre-clearing step was performed by incubating the whole cell lysates with 25 µl beads for 2 h at 4°C. The beads were then discarded, and the cell lysates were incubated with antibody overnight as already described.

### Protein expression, purification, and labeling

All proteins used in this study were overexpressed in Rosetta (DE3, pLysS) *Escherichia coli* (MERCK) cells using 0.4 mM isopropyl 1-thio-D-galactopyranoside (Inalco Pharmaceuticals) induction overnight at 16 °C. Purification of proteins was performed on Ni Sepharose 6 Fast Flow sepharose (Cytiva) in 50 mM HEPES (pH 7.5), 300 mM NaCl, 1 mM TCEP, 10 mM imidazole and 10% (w/v) glycerol and eluted using 250 mM imidazole (pH 7.5) in the same buffer. UBA6 and E2 variants were further desalted using PD-10 Columns (Cytiva) into Loading buffer containing 20 mM HEPES (pH 7.5), 150mM NaCl, and 10% (w/v) glycerol. Concentrated protein aliquots were stored at 80 °C. All protein concentrations indicated correspond to total protein and are based on UV absorbance at 280 nm.

Fluorescein-5-Maleimide (AnaSpec) was attached to ubiquitin following the directions as previously described ^40^. Briefly, proteins in 20mM Tris (pH 7.5), 150mM NaCl and 1mM TCEP were incubated for 2 hours in the presence of the fluorophore at room temperature such that the label : protein ratio would be 4. To quench the reaction, beta-mercaptoethanol was added at a ratio of 10:1 to fluorescein. Fluorescein-labeled ubiquitin was then separated from free dye on PD10 desalting columns (Cytiva) and was stored at 80 °C.

### E1 and E2 loading assays

All loading assays were performed at 32 °C in a buffer containing 20 mM HEPES (pH 7.5), 150 mM NaCl and 10% (w/v) glycerol. E1 and E2 loading assays were performed using 3 µM fluorescein ubiquitin, 10 nM to 1 µM E1, 1 µM E2, and 2.5mM concentrations each of ATP and MgCl_2_. Reactions were stopped using nonreducing SDS-PAGE loading buffer. Samples were separated on 4–20% Tris-Glycine NuPAGE gels (Thermo) in Tris-Glycine buffer. Band detection in E1 and E2 loading assays was performed using an Alliance Q9 imager (Uvitec Cambridge). The different qualitative end-point assays were performed using freshly thawed protein aliquots, and the results obtained were reproducible across at least three different protein batches. All constructs and conditions were carried out in triplicate.

### *In silico* structural analysis

We constructed models for UBA6 using the Protein Homology/analogY Recognition Engine V 2.0 (PHYRE2) and the AlphaFold server ^43, 44^. Further idealization of the geometry was achieved by five cycles of minimization with Refmac5. The model of full length USE1 was downloaded from the AlphaFold server. Structural alignment of USE1 and UBA6 on Ubc4:UBA1:Ub complex (PDB code 4II2) ^45^ provided a structural model for the USE1:UBA6 complex. Structure visualization and figures preparations were performed with PyMOL (Molecular Graphics System, http://www.pymol.org). We employed the continuum solvation method APBS (Adaptive Poisson– Boltzmann Solver) with the CHARMM force field to calculate the electro-potential surface of UBA1 and UBA6 ^46^.

### Confocal microscopy

The cells were grown on coverslips, and then washed and fixed in 4% Paraformaldehyde for 10-15 minutes before being permeabilized with 0.1% Triton X-100. A solution of, 1% BSA in PBS was used to block both primary and secondary antibodies. The primary antibody was added at a ratio of 1:100 and incubated for at least 1 h at room temperature while the secondary antibody (1:300 Invitrogen) was allowed to incubate with the sample for 30 min at room temperature. Neurons were permeabilized in 2% BSA + 0.1% Triton, and blocked with 2% BSA and the primary antibodies were incubated overnight at 4°C, at a ratio of 1:150, with the secondary antibody incubated for 2 h at room temperature at 1:500. A Zeiss 710 confocal microscope was utilized for confocal imaging with a 63X oil-immersion lens. Nuclear staining was detected by staining with DRAQ5. For quantification, the operator was blinded to the outcome of the experiment when selecting suitably similar fields to image for subsequent computerized analysis. For the colocalization experiments, the association of PHOX2B and UBA6 outside the nucleus was measured by selecting PHOX2B positive cells manually using Fiji, and excluding the nuclei by segmenting and removing the nuclear channel. The colocalization between the channels of interest was then measured using the JACops plugin in Fiji with the default parameters, and Pearson’s correlation coefficient was calculated. The cytoplasmic intensity was measured for the channel of interest and the mean gray value was recorded after excluding the nuclei as already described. For the localization experiments related to UBA6 in neurons, the association of UBA6 with the cell body was examined by selecting the neuronal cell bodies manually using Fiji ^47^. For the localization of UBA6 with neurites, the cell bodies were selected manually and removed. Arc intensity in the neurons was measured by circulating the neuronal cell body together with the first neurite junction manually using Fiji. The mean integrated value and the area for each neuron were recorded. The value of the mean divided by the area was used for statistical analysis.

### Bioinformatics analysis

USE1/UBE2Z human homologs were searched against the Uniprot (ref pubmed id 29425356) and NCBI databases using BLAST (ref pubmed id 2231712). Prosite (ref pubmed id 23161676) was used to scan for alanine residues motifs with between 6 and 10 continuous alanine residues in the BLAST results (a search for proteins containing polyalanine stretches in the ubiquitin cascades). Alignments of the E2 family in vertebrates and across all databases were calculated using MAFFT (ref pubmed id 28968734). The figures were generated using Jalview (ref pubmed id 19151095).

### Statistics

Basic data handling was performed in Microsoft Excel. For single comparisons, the statistical significance of the difference between experimental groups was determined using two-tailed Student’s t-test with the Prism GraphPad software v.9. Comparisons of multiple means were made by one-way or two-way analysis of variance (ANOVA) followed by the Tukey’s, Dunnett’s and Sidak’s post hoc tests to determine statistical significance. Differences were considered statistically significant for *P* < 0.05. Sample sizes were chosen based on extensive experience with the assays we have performed. The experiments were appropriately randomized. For primary neurons transduced with lentiviral vectors, we used independent cultures prepared from brains of mouse embryos taken from different females. For iPSC-derived neurons, three independent cultures from different differentiation days were considered for analysis, and for cell-line based experiments, we considered replicates performed in different days. Errors bars shown in the figures are standard errors of the mean (s.e.m).

## Data availability

Available protein structures were from PDB code 4II2, 6DC6, 1Y8Q, 2NVU. Structural analysis was supported by AlphaFold Protein Structure Database, protein sequences were from UniProt.

## Acknowledgements

We are grateful for funding from Azrieli Foundation (grant to A.A), CCHS Foundation/CCHS Network (grant to A.A and G.D.V), Israel Science Foundation (ISF) 1621/18 and 2327/18 (grants to G.D.V), and the ISF grant 1440/21 to G.P. Yoran Institute for Human Genome Research (scholarship to F.A.S). We thank Yad Laneshima for assistance in patient recruitment; A. Yeheskel and H. Benyamini for bioinformatics analysis; D. Fornasari, D.C Rubinsztein, A. Aichem, M. Groettrup, Y. Tokita, and D. Hnisz for contributing reagents.

## Contributions

A.A developed the study rationale. A.A, G.D.V and G.P wrote the manuscript with inputs from all authors. F.A.S performed and analyzed cell line and primary neuron experiments and was involved in all other experiments of the study. G.D.V and D.F designed iPSC experiments, D.F performed the experiments involving iPSCs, Y.B performed the molecular biology and in vitro ubiquitination experiments with assistance from A.K, G.P generated and analyzed the structural models. A.A. supervised the study.

---

**Extended Data Fig. 1.**
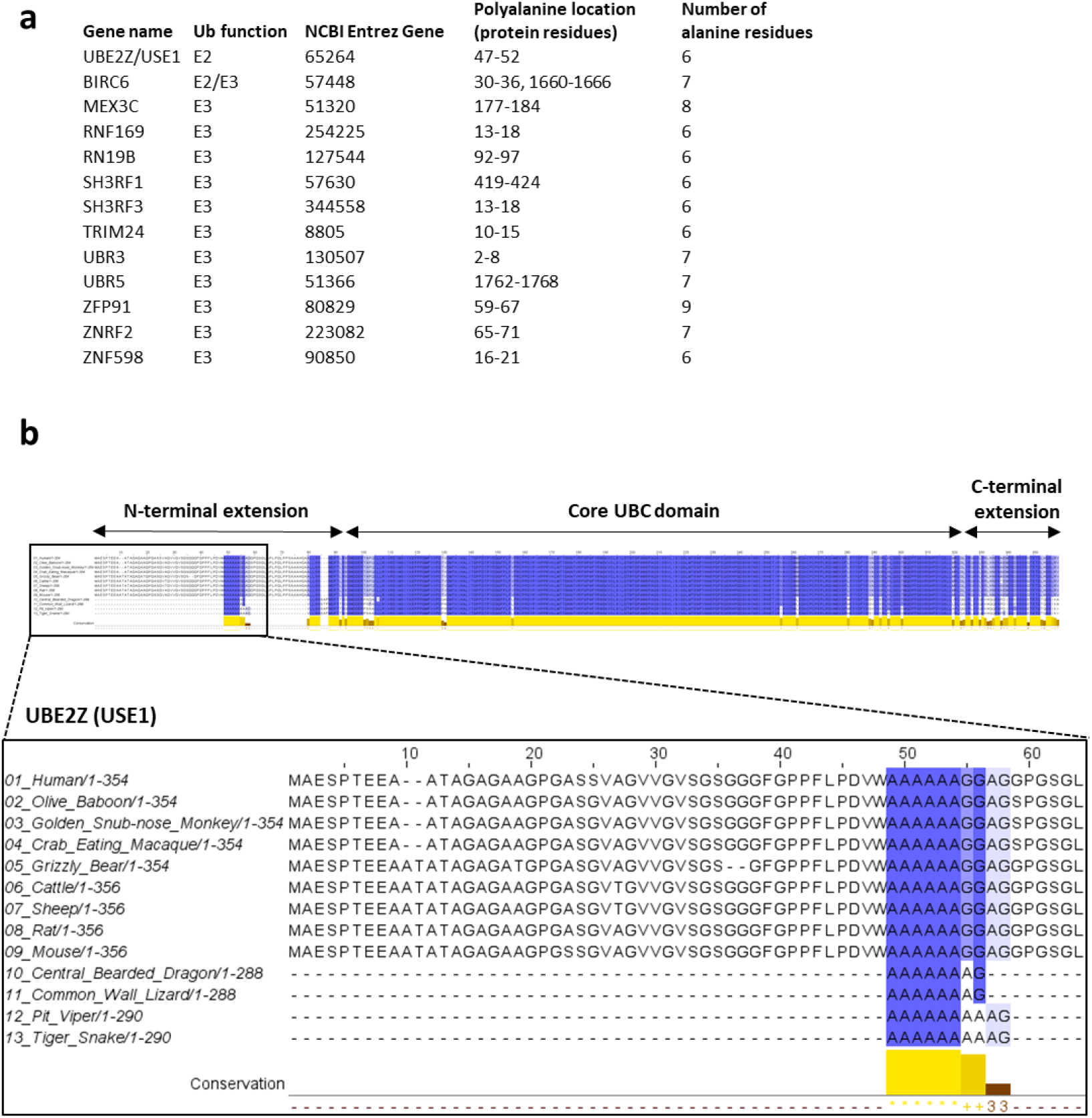
Analysis of alanine repeats in the ubiquitin system. **a,** Analysis of alanine repeats domains in the human ubiquitin cascades comprising E1, E2, and E3 enzymes. **b,** A multiple sequence alignment of USE1 homologs from different vertebrates. The alignment is colored according to sequence identity including the N-terminus containing the polyalanine stretch.

**Extended Data Fig. 2.**
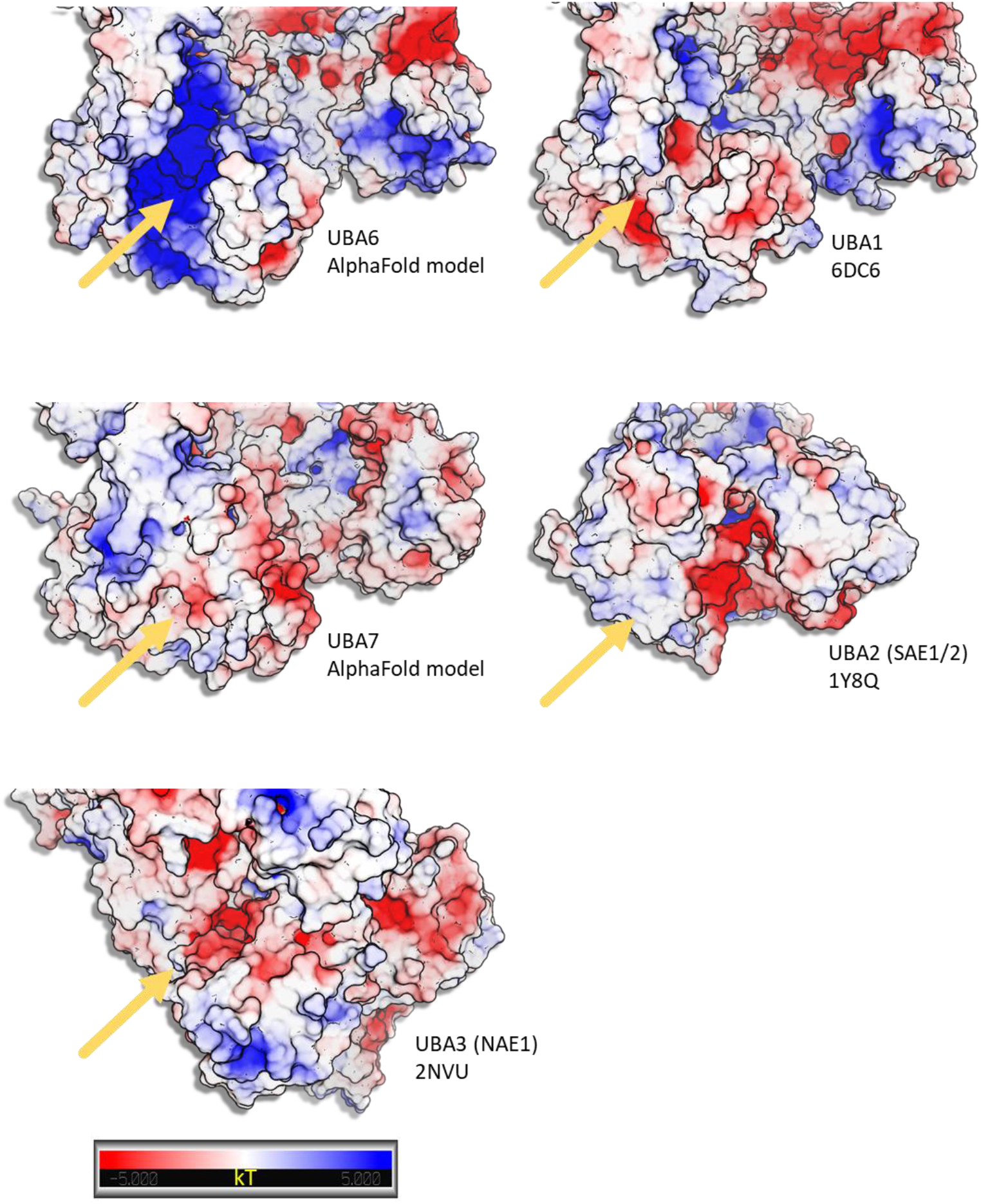
Structures and models of the SCCH domains of the cannonical E1 ubiquitin-like activating enzymes. AlphaFold models of UBA6 and UBA7 and the crystal structures of UBA1, UBA2 and UBA3 are shown. The structures were aligned and electrostatic potential was clulated as descibed in the methods. The yellow arrows indicate the location of the groove within the SCCH domains. UBA6, UBA1 and UBA7 form an extended lobe within the SCCH, which is missing in UBA2 and UBA3. The groove in UBA7 is covered and do not exist in UBA2 and UBA3. The grooves in UBA1 and UBA6 are highly similar in terms of structure but present significantly different electro potential surfaces. The gradiant from negative (red) to positve (blue) charge is shown. The figure was prepared by PyMol.

**Extended Data Fig. 3.**
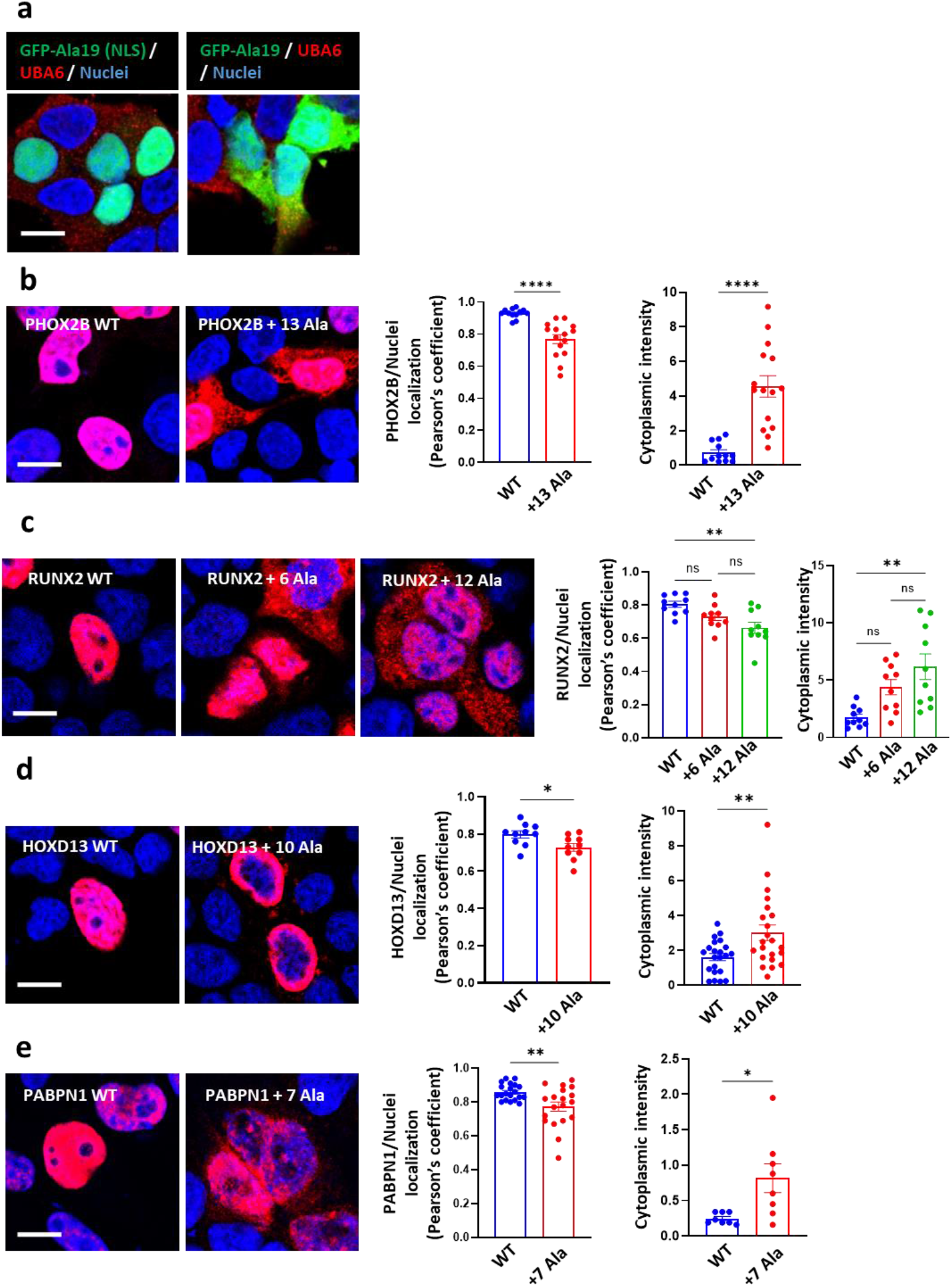
Polyalanine expansion mutations cause cytoplasmic mislocalization of different nuclear proteins. HEK293T cells were transfected with the indicated constructs, and were subjected to immunostaining. **a**, GFP-19Ala with nuclear localization sequence (NLS) or GFP-19Ala, labeled for endogenous UBA6. **b**, HA-PHOX2B WT and HA-PHOX2B +13Ala. **c**, HA-RUNX2 WT, HA-RUNX2 +6Ala, and HA-RUNX2+ 12Ala. **d**, HA-HOXD13 WT and HA-HOXD13 +10Ala. **e**, HA-PABPN1 WT and HA-PABPN1 +7Ala. Image scale bar 20 μm. Quantification of the association of HA-tagged proteins (labeled in red) with the nucleus (labeled in blue, Pearson’s coefficient) is indicated as well as the cytoplasmic intensity. Results are the average values from cells in different imaged fields. Between 60-100 transfected cells were analyzed. Results are mean ± s.e.m. Unpaired 2-tailed t-test (**b,d,e)** and one-way ANOVA Tukey’s test (**c**). ns non-significant, \**P* < 0.05, \*\**P* < 0.01, \*\*\*\**P* < 0.0001.

**Extended Data Fig. 4.**
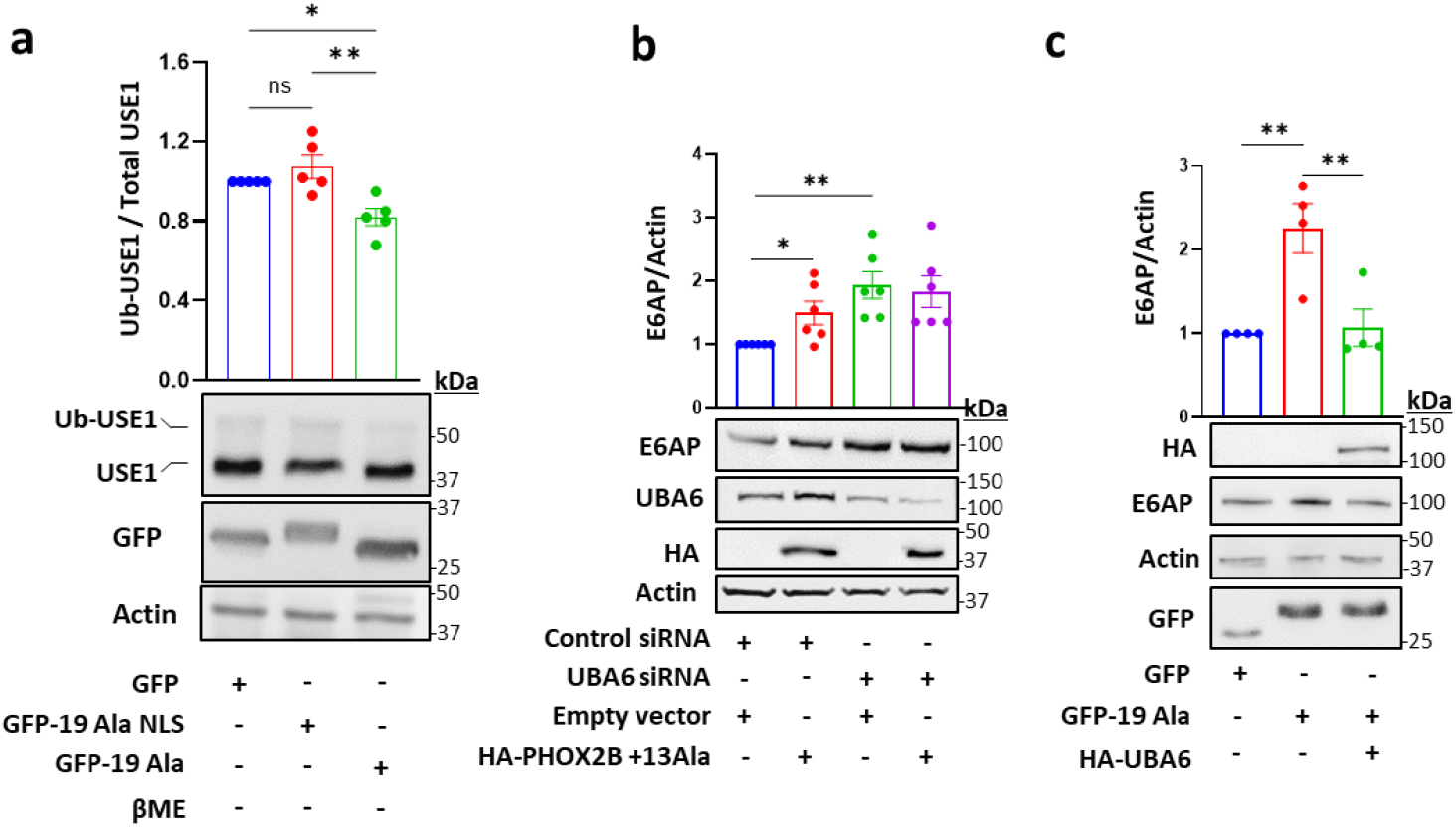
Isolated polyalanine stretches regulate USE1 ubiquitin loading and E6AP levels. **a**, HEK293T cells were transfected with empty GFP, GFP-polyAla, and GFP-polyAla with a nuclear localization sequence (NLS). Endogenous USE1 ubiquitin loading was analyzed in βME-untreated cell lysates. **b**, Control and UBA6-depleted HEK293T cells were transfected with empty vector or mutant PHOX2B and analyzed for E6AP levels. **c,** Cells were transfected with empty GFP, or GFP-polyAla with or without HA-UBA6, and analyzed for E6AP levels. Results are mean ± s.e.m. (**a**) One-way ANOVA Tukey’s test, n=5 (**b**) Unpaired 2-tailed t-test, n=6. (**c**) One-way ANOVA, Tukey’s test, n=4. n.s not significant, \**P* < 0.05, \*\**P* < 0.01.

**Extended Data Fig. 5.**
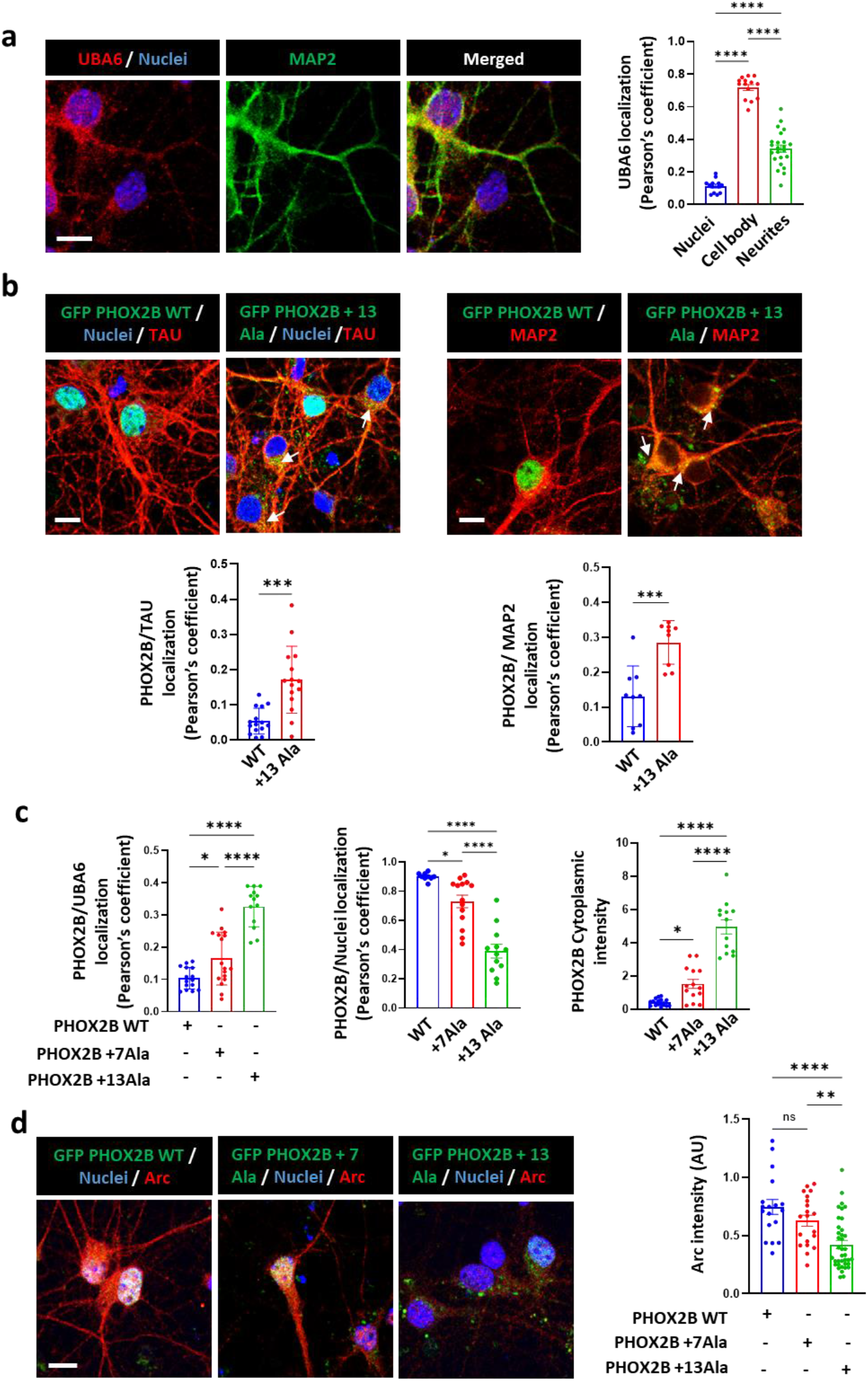
Analysis of UBA6 and mutant PHOX2B association and effects on Arc levels in primary neurons. **a,** Mouse primary cortical neurons were labeled for endogenous UBA6 and MAP2. Quantification of the association of UBA6 with the neuronal nucleus, cell body, and neurites is shown (Pearson’s coefficient). **b**, Mouse primary cortical neurons were transduced with lentiviral vectors expressing GFP-tagged WT PHOX2B or mutant PHOX2B (+13Ala), and were labeled for endogenous MAP2 and TAU (non-nuclear fraction of mutant PHOX2B marked with arrows). The quantification of the association of GFP-PHOX2B with the nucleus, MAP2, and TAU is shown (Pearson’s coefficient), as well as GFP-PHOX2B cytoplasmic intensity. **c**. Quantification of UBA6 association with WT and mutant (+7 Ala, +13 Ala) GFP-PHOX2B (Pearson’s coefficient) related to Fig. 3g. **d,** The primary neurons were transduced with the GFP-PHOX2B lentiviral vectors and were labeled for Arc. The quantification of Arc intensity in the GFP-PHOX2B expressing neurons is presented. Image scale bar 10 μm. Results are mean ± s.e.m representing the average values from neurons in different imaged fields. (**a**) **\***\*\*\**P* < 0.0001 One– way ANOVA Tukey’s test. n= 50 neurons analyzed. (**b**) **\*\*\*** *P* < 0.001 unpaired 2-tailed t-test, n=30-50 neurons analyzed for WT PHOX2B and mutant PHOX2B +13Ala. (**c**) **\****P* < 0.05, **\***\*\*\**P* < 0.0001 One–way ANOVA Tukey’s test. n=30, n=60, and n=90 neurons analyzed for WT PHOX2B, mutant PHOX2B +7Ala and +13Ala, respectively. (**d**) ns non-significant, \*\**P* < 0.01, \*\*\*\**P* < 0.0001, one-way ANOVA Tukey’s test. n=20, n=30 and n=100 neurons analyzed for WT PHOX2B, mutant PHOX2B +7Ala and +13Ala, respectively.

**Extended Data Fig. 6.**
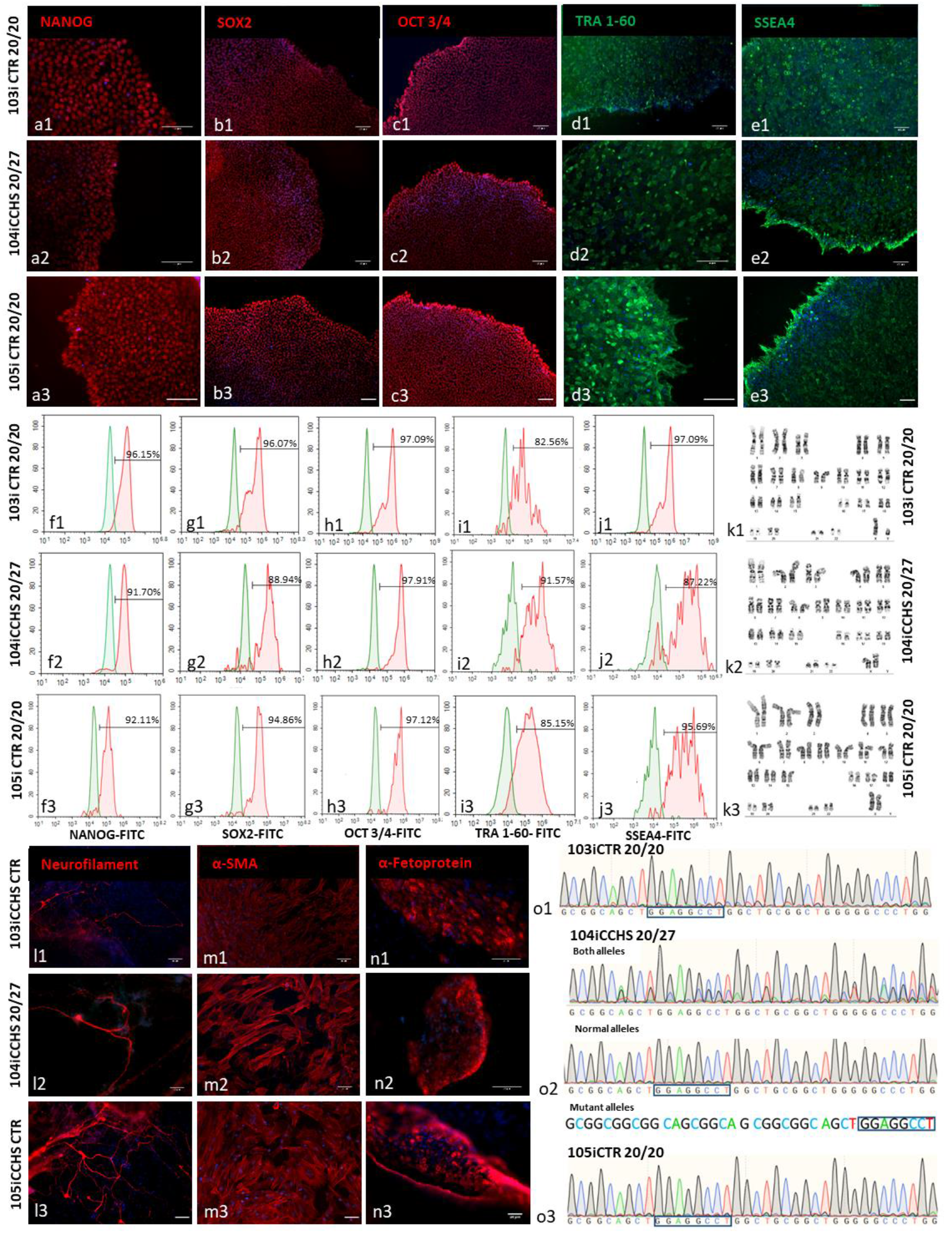
Generation and characterization of CCHS patient and family relative-derived iPSCs. Skin punch biopsies were collected from a 2-year-old female patient with CCHS who harbors a heterozygous 27 polyalanine expansion in PHOX2B (104iCCHS 20/27), and from her healthy sister (105iCTR 20/20). A biopsy was also collected from the healthy father (103iCTR 20/20) of the 4-year-old male patients who harbors a heterozygous 25 polyalanine expansion in PHOX2B (102iCCHS 20/25). Patient-specific fibroblasts were electroporated with non-integrating reprogramming episomal plasmids. **a-e**, Immunocytochemistry for pluripotency markers (NANOG, SOX2, OCT3/4 TRA-1-60, SSEA4). **f-j**, Flow cytometry analysis for pluripotency markers (red-NANOG, SOX2, OCT3/4 TRA-1-60, SSEA4). **k**, Karyotype. **l-n**, Embryoid bodies (EBs) were generated and allowed to spontaneously differentiate for 21 days. Differentiated EBs express the ectoderm marker neurofilament, the mesoderm marker α-SMA, and the endoderm marker α-fetoprotein. **o**, Sequencing of the 3rd PHOX2B exon confirms the heterozygous +7 polyalanine expansion in the CCHS patient, but not in the healthy controls. Image scale bar 100 µm.

**Extended Data Fig. 7.**
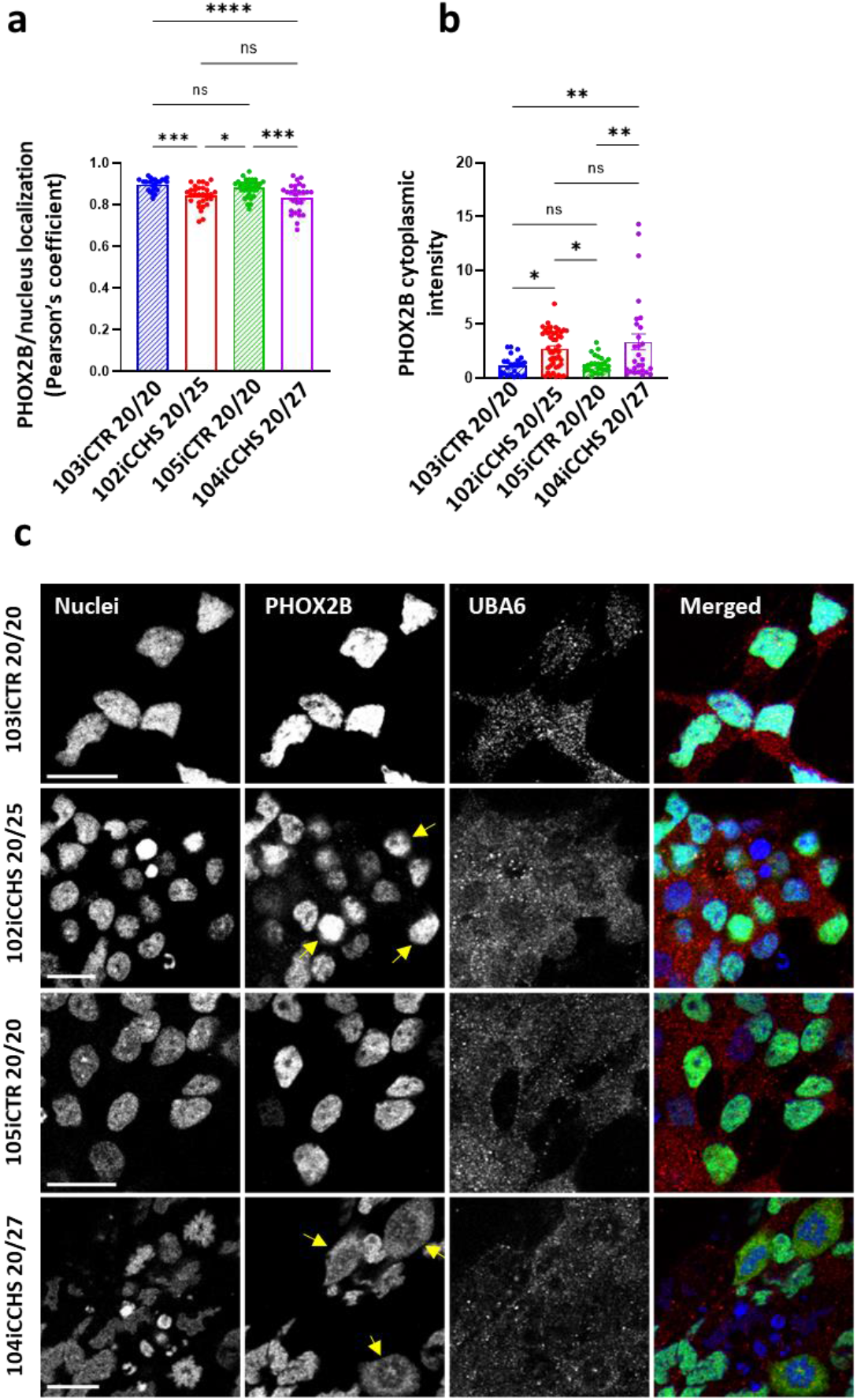
PHOX2B cytoplasmic mislocalization in patient-derived autonomic neurons. **a-b**, Quantification of the association of endogenous PHOX2B with the nucleus (Pearson’s coefficient) in autonomic neurons from control and CCHS patients. Quantification is shown also for PHOX2B cytoplasmic intensity. Results are mean ± s.e.m. \**P* < 0.05, \*\**P* < 0.01, \*\*\**P* < 0.001 \*\*\*\**P* < 0.0001 one-way ANOVA Tukey’s test. The results represent additional analysis from the same neurons analyzed in Fig. 4b. **c**, iPSC-derived human autonomic neurons from control and CCHS patients were labeled by nuclear staining (colored blue), and for endogenous PHOX2B (colored green) and endogenous UBA6 (colored red). Images indicate events of severe cytoplasmic mislocalization of PHOX2B (marked with arrows). Scale bar 10 μm.

## References

1. Amiel, J. et al. Polyalanine expansion and frameshift mutations of the paired-like homeobox gene PHOX2B in congenital central hypoventilation syndrome. Nat Genet 33, 459–461 (2003).

2. Muragaki, Y., Mundlos, S., Upton, J. & Olsen, B.R. Altered growth and branching patterns in synpolydactyly caused by mutations in HOXD13. Science 272, 548–551 (1996).

3. Mundlos, S. et al. Mutations involving the transcription factor CBFA1 cause cleidocranial dysplasia. Cell 89, 773–779 (1997).

4. Laumonnier, F. et al. Transcription factor SOX3 is involved in X-linked mental retardation with growth hormone deficiency. Am J Hum Genet 71, 1450–1455 (2002).

5. Richard, P. et al. Correlation between PABPN1 genotype and disease severity in oculopharyngeal muscular dystrophy. Neurology 88, 359–365 (2017).

6. Brown, L.Y. et al. Holoprosencephaly due to mutations in ZIC2: alanine tract expansion mutations may be caused by parental somatic recombination. Hum Mol Genet 10, 791–796 (2001).

7. Jin, J., Li, X., Gygi, S.P. & Harper, J.W. Dual E1 activation systems for ubiquitin differentially regulate E2 enzyme charging. Nature 447, 1135–1138 (2007).

8. Pelzer, C. et al. UBE1L2, a novel E1 enzyme specific for ubiquitin. J Biol Chem 282, 23010–23014 (2007).

9. Usdin, K. The biological effects of simple tandem repeats: lessons from the repeat expansion diseases. Genome Res 18, 1011–1019 (2008).

10. Albrecht, A. & Mundlos, S. The other trinucleotide repeat: polyalanine expansion disorders. Curr Opin Genet Dev 15, 285–293 (2005).

11. Amiel, J., Trochet, D., Clement-Ziza, M., Munnich, A. & Lyonnet, S. Polyalanine expansions in human. Hum Mol Genet 13 Spec No 2, R235–243 (2004).

12. Amer-Sarsour, F., Kordonsky, A., Berdichevsky, Y., Prag, G. & Ashkenazi, A. Deubiquitylating enzymes in neuronal health and disease. Cell Death Dis 12, 120 (2021).

13. Lee, P.C. et al. Altered social behavior and neuronal development in mice lacking the Uba6-Use1 ubiquitin transfer system. Mol Cell 50, 172–184 (2013).

14. Matsuura, T. et al. De novo truncating mutations in E6-AP ubiquitin-protein ligase gene (UBE3A) in Angelman syndrome. Nat Genet 15, 74–77 (1997).

15. Ye, Y. & Rape, M. Building ubiquitin chains: E2 enzymes at work. Nat Rev Mol Cell Biol 10, 755–764 (2009).

16. Schelpe, J., Monte, D., Dewitte, F., Sixma, T.K. & Rucktooa, P. Structure of UBE2Z Enzyme Provides Functional Insight into Specificity in the FAT10 Protein Conjugation Machinery. J Biol Chem 291, 630–639 (2016).

17. Bohnsack, R.N. & Haas, A.L. Conservation in the mechanism of Nedd8 activation by the human AppBp1-Uba3 heterodimer. J Biol Chem 278, 26823–26830 (2003).

18. Whitby, F.G., Xia, G., Pickart, C.M. & Hill, C.P. Crystal structure of the human ubiquitin-like protein NEDD8 and interactions with ubiquitin pathway enzymes. J Biol Chem 273, 34983–34991 (1998).

19. Schulman, B.A. & Harper, J.W. Ubiquitin-like protein activation by E1 enzymes: the apex for downstream signalling pathways. Nat Rev Mol Cell Biol 10, 319–331 (2009).

20. Lee, I. & Schindelin, H. Structural insights into E1-catalyzed ubiquitin activation and transfer to conjugating enzymes. Cell 134, 268–278 (2008).

21. Huang, D.T. et al. Basis for a ubiquitin-like protein thioester switch toggling E1-E2 affinity. Nature 445, 394–398 (2007).

22. Lois, L.M. & Lima, C.D. Structures of the SUMO E1 provide mechanistic insights into SUMO activation and E2 recruitment to E1. EMBO J 24, 439–451 (2005).

23. Akimoto, G., Fernandes, A.P. & Bode, J.W. Site-Specific Protein Ubiquitylation Using an Engineered, Chimeric E1 Activating Enzyme and E2 SUMO Conjugating Enzyme Ubc9. ACS Cent Sci 8, 275–281 (2022).

24. Kishino, T., Lalande, M. & Wagstaff, J. UBE3A/E6-AP mutations cause Angelman syndrome. Nat Genet 15, 70–73 (1997).

25. Glessner, J.T. et al. Autism genome-wide copy number variation reveals ubiquitin and neuronal genes. Nature 459, 569–573 (2009).

26. Greer, P.L. et al. The Angelman Syndrome protein Ube3A regulates synapse development by ubiquitinating arc. Cell 140, 704–716 (2010).

27. Kuhnle, S., Mothes, B., Matentzoglu, K. & Scheffner, M. Role of the ubiquitin ligase E6AP/UBE3A in controlling levels of the synaptic protein Arc. Proc Natl Acad Sci U S A 110, 8888–8893 (2013).

28. Dauger, S. et al. Phox2b controls the development of peripheral chemoreceptors and afferent visceral pathways. Development 130, 6635–6642 (2003).

29. Pattyn, A., Morin, X., Cremer, H., Goridis, C. & Brunet, J.F. The homeobox gene Phox2b is essential for the development of autonomic neural crest derivatives. Nature 399, 366–370 (1999).

30. Vanderlaan, M., Holbrook, C.R., Wang, M., Tuell, A. & Gozal, D. Epidemiologic survey of 196 patients with congenital central hypoventilation syndrome. Pediatr Pulmonol 37, 217–229 (2004).

31. Trang, H. et al. Proceedings of the fourth international conference on central hypoventilation. Orphanet J Rare Dis 9, 194 (2014).

32. Falik, D. et al. Generation and characterization of iPSC lines (BGUi004-A, BGUi005-A) from two identical twins with polyalanine expansion in the paired-like homeobox 2B (PHOX2B) gene. Stem Cell Res 48, 101955 (2020).

33. Ashkenazi, A. et al. Polyglutamine tracts regulate beclin 1-dependent autophagy. Nature 545, 108–111 (2017).

34. Paulson, H.L., Shakkottai, V.G., Clark, H.B. & Orr, H.T. Polyglutamine spinocerebellar ataxias – from genes to potential treatments. Nat Rev Neurosci 18, 613–626 (2017).

35. Dubreuil, V. et al. A human mutation in Phox2b causes lack of CO_2_ chemosensitivity, fatal central apnea, and specific loss of parafacial neurons. Proc Natl Acad Sci U S A 105, 1067–1072 (2008).

36. Ramanantsoa, N. et al. Ventilatory response to hyperoxia in newborn mice heterozygous for the transcription factor Phox2b. Am J Physiol Regul Integr Comp Physiol 290, R1691–1696 (2006).

37. Basu, S. et al. Unblending of Transcriptional Condensates in Human Repeat Expansion Disease. Cell 181, 1062–1079 e1030 (2020).

38. Chen, Z. et al. Down-regulation of UBA6 exacerbates brain injury by inhibiting the activation of Notch signaling pathway to promote cerebral cell apoptosis in rat acute cerebral infarction model. Mol Cell Probes 53, 101612 (2020).

39. Shilling, P.J. et al. Improved designs for pET expression plasmids increase protein production yield in Escherichia coli. Commun Biol 3, 214 (2020).

40. Keren-Kaplan, T. et al. Synthetic biology approach to reconstituting the ubiquitylation cascade in bacteria. EMBO J 31, 378–390 (2012).

41. Kamitani, T., Kito, K., Nguyen, H.P. & Yeh, E.T. Characterization of NEDD8, a developmentally down-regulated ubiquitin-like protein. J Biol Chem 272, 28557–28562 (1997).

42. Kirino, K., Nakahata, T., Taguchi, T. & Saito, M.K. Efficient derivation of sympathetic neurons from human pluripotent stem cells with a defined condition. Sci Rep 8, 12865 (2018).

43. Kelley, L.A., Mezulis, S., Yates, C.M., Wass, M.N. & Sternberg, M.J. The Phyre2 web portal for protein modeling, prediction and analysis. Nat Protoc 10, 845–858 (2015).

44. Jumper, J. et al. Highly accurate protein structure prediction with AlphaFold. Nature 596, 583–589 (2021).

45. Olsen, S.K. & Lima, C.D. Structure of a ubiquitin E1-E2 complex: insights to E1-E2 thioester transfer. Mol Cell 49, 884–896 (2013).

46. Baker, N.A., Sept, D., Joseph, S., Holst, M.J. & McCammon, J.A. Electrostatics of nanosystems: application to microtubules and the ribosome. Proc Natl Acad Sci U S A 98, 10037–10041 (2001).

47. Schindelin, J. et al. Fiji: an open-source platform for biological-image analysis. Nat Methods 9, 676–682 (2012).

